# Environmental filtering and host identity collectively shape root-associated microbiomes of Ericaceae and ectomycorrhizal plants in fumarole fields

**DOI:** 10.64898/2026.09.25.754363

**Authors:** Akifumi Murata, Mikihito Noguchi, Ibuki Hayashi, Keitaro Fukushima, Hirokazu Toju

## Abstract

**Background:** Symbiosis with microbes is a key strategy that has enabled plants to colonize extreme environments. Since the benefits conferred by root-associated microbes depend on both environmental conditions and host-microbe combinations, plant adaptation to harsh environments is closely linked to the assembly of root microbial communities. Understanding how environmental and host filtering jointly shape these communities is therefore fundamental to elucidating the mechanisms underlying plant adaptation to extreme environments.

**Results:** In this study, we investigated the differentiation of root-associated prokaryotic and fungal communities and individual operational taxonomic units (OTUs) across two contrasting habitats surrounding fumaroles, solfatara-field and forest-edge habitats, and six dominant Ericaceae and ectomycorrhizal plant taxa. Prokaryotic and fungal OTUs rarely exhibited strong preferences for both habitat and host identity. Instead, many of prokaryotic and fungal OTUs specialized to one of these niches, collectively generating root microbial communities differentiated by both factors. Nonetheless, striking specializations in habitat and host niches were observed in the fungal family Hyaloscyphaceae (Helotiales). To gain insight into the evolutionary basis of microbial specialization, we examined phylogenetic signals in preference phenotypes. The resulting weak phylogenetic signals in these preference phenotypes further suggest that this fungal clade has undergone substantial ecological divergence.

**Conclusion:** Overall, our findings indicate that root-associated microbial communities in extreme environments are assembled through the accumulation of microbial taxa specialized to either habitat or host, and that strong ecological specialization in fungi can arise with little phylogenetic constraint.

## Background

Symbiosis with bacteria and fungi is a crucial adaptive strategy that supports plant establishment and growth in extreme environments [1–7]. Plant roots serve as a primary site of interactions with diverse soil-borne microbes, and the microbial community formed in the root endosphere can significantly influence host fitness [2, 8–12]. In extreme environments, stressors such as salinity and low temperatures act as potent filters that determine the composition of soil microbial communities and the potential microbes to colonize the root endosphere [13–15]. Some of the root endophytes selected under these environmental conditions can confer habitat-specific stress tolerance to their host (“habitat-adapted symbiosis”) [6]. However, because of compatibility constraints (interaction specificity) between plant identities and microbial strains [16–18], microbes selected by harsh environmental conditions do not necessarily succeed in colonizing the root endosphere. They undergo further filtering by the host plant during the transition from soil to root [19–21]. Therefore, to understand plant adaptation to extreme environments mediated by microbial symbiosis, it is essential to elucidate how environmental and host filtering processes select microbes and shape root-associated microbial communities.

To understand how root-associated microbial communities are assembled, a bottom-up perspective is required, in which community-level compositional differences are viewed as the collective outcome of heterogeneous niche responses among individual microbial taxa. Previous studies have largely examined how root-associated microbial communities differ in response to environmental and host factors, highlighting the importance of these factors in community assembly [19, 20, 22–24]. However, individual microbial taxa can vary widely in their responses to environmental conditions, host identity, and their interactions [25–28]. Thus, community-level analyses alone cannot identify which taxon-specific niche responses collectively drive the compositional differences observed across environments and hosts. To characterize such niche differentiation among individual microbial taxa, it is necessary to disentangle the effects of environmental conditions and host identity on microbial taxon distributions. However, in natural ecosystems, host plant composition often varies along environmental gradients, causing environmental and host effects to become confounded [29, 30]. This confounding makes it difficult to determine whether individual microbial taxa are associated primarily with environmental conditions, host identity, or their interaction.

Fumarole fields provide a suitable natural system for addressing this challenge. These areas are characterized by multiple environmental stresses, including high temperature, strongly acidic soils, toxic gases, and nutrient limitation [31], with soil conditions varying sharply over distances of only a few meters [32]. Despite these steep environmental gradients, Ericaceae shrubs and ectomycorrhizal plants such as Pinaceae and Betulaceae coexist and are widely distributed across contrasting habitats. Although these plants often form sympatric patches and share some root-associated endophytes [33], they also harbor host-specific root fungal communities [34, 35]. This combination of steep environmental gradients and shared host species with distinct root endophyte associations provides an ideal natural system for disentangling environmental and host filtering.

In this study, we investigated how environmental and host filtering shape root-associated microbial communities across contrasting habitats in fumarole fields, focusing on how OTU-level niche differentiation underlies community-level differentiation. We conducted field surveys in two fumarole fields in Japan, targeting four Ericaceae and two ectomycorrhizal plant genera that commonly occur across two contrasting habitats: solfatara-field habitats, characterized by sparsely vegetated bare ground near fumaroles, and forest-edge habitats. Root-associated prokaryotic and fungal communities were characterized using high-throughput DNA metabarcoding. We hypothesized that strong abiotic filtering and host-specific recruitment promote the colonization of habitat- and host-specialized microbial OTUs, thereby driving community differentiation across contrasting habitats and host identities. To test this hypothesis, we first quantified the relative contributions of habitat and host plant identity to community differentiation. We then identified the habitat and host preferences of individual microbial OTUs to determine how OTU-level niche differentiation contributes to community-level patterns. We further examined whether these ecological preferences are phylogenetically conserved, providing additional insight into the evolutionary constraints underlying microbial specialization. Together, these analyses clarify how root microbiome structure is organized across steep abiotic gradients and multiple host species in extreme environments, providing insights into how root-associated microbiomes may contribute to plant persistence under harsh conditions.

## Methods

### Study site and sampling

Sampling was conducted from June 5 to June 6, 2024, in the Onikobe volcanic region, located at the southern foot of Mt. Arao, Miyagi Prefecture, Japan. We surveyed two fumarole fields: Arayu-Jigoku (38.82 °N, 140.73 °E), at approximately 600 m a.s.l., and Ofukasawa (38.81 °N, 140.71°E), at approximately 500 m a.s.l. The surveyed areas were approximately 300 m × 20 m in Arayu-Jigoku and 200 m × 20 m in Ofukasawa.

Numerous fumaroles emit high-temperature water vapor and sulfur-containing gases along the valleys in these solfatara fields, forming sparsely vegetated bare areas (Fig. 1A). Despite these harsh conditions, small patches of Ericaceae shrubs, such as *Rhododendron multiflorum* and *Eubotryoides grayana*, occur near the fumaroles (hereafter, solfatara-field habitat; Fig. 1A). Outside these bare areas, more densely vegetated forest-edge habitats are established by shrubs and trees, primarily Ericaceae and Pinaceae, together with some Betulaceae and Fagaceae species (hereafter, forest-edge habitat). Solfatara field soil is generally characterized by the depletion of soluble base cations and soluble metal ions [36]. Forest soil developed near such a solfatara field may represent a more moderate environment in terms of the availability of cations required for plant growth. However, increased solubilization of aluminum by organic acids released from plant roots in the forest environment can promote hydrolysis reactions that release additional protons, further lowering the already low pH [37]. Thus, the solfatara-field and the adjacent forest-edge zone constitute harsh environments for plant growth, but for different reasons.

**Fig 1.**
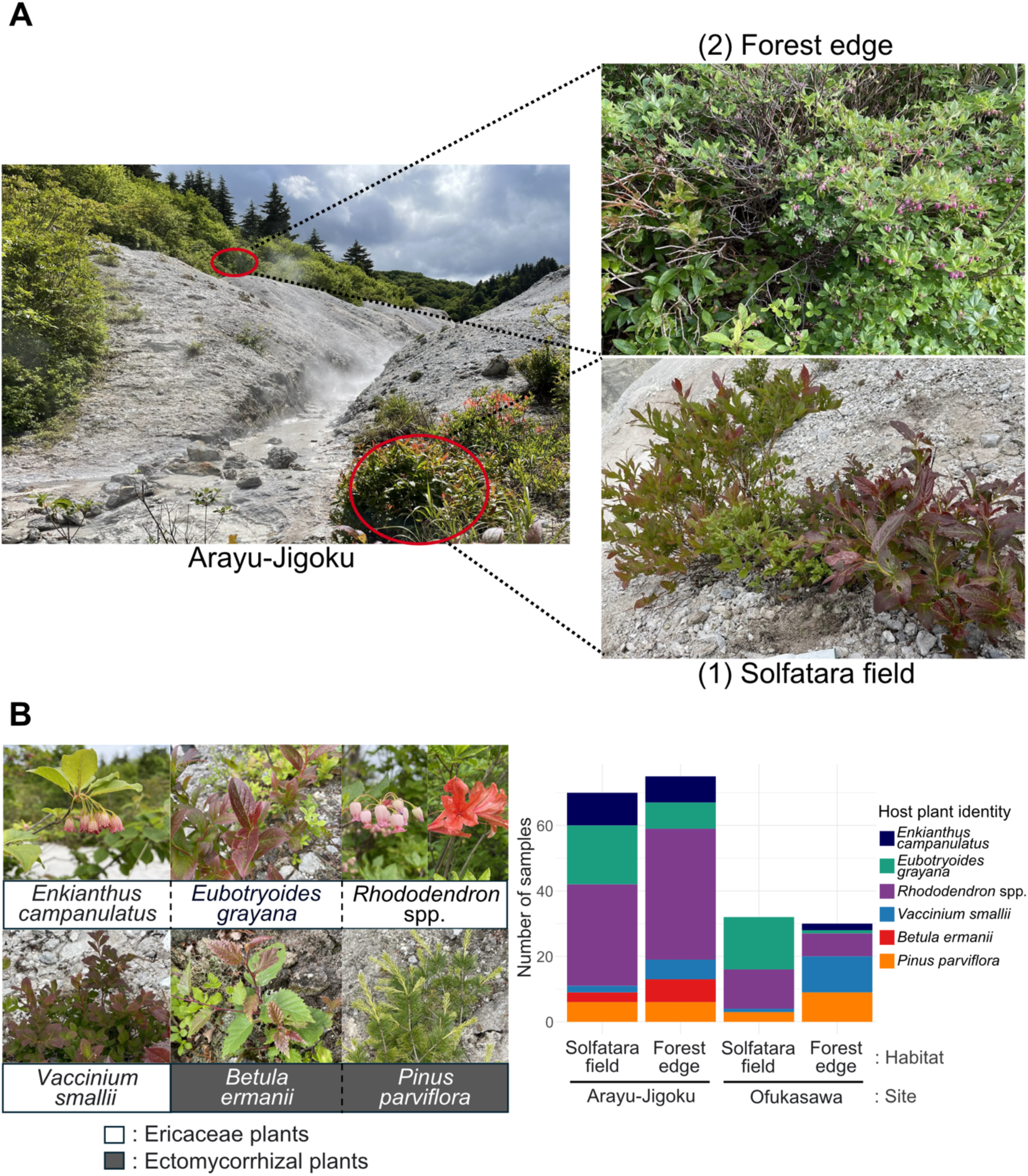
Overview of the study sites and target plant samples. (A) Photograph of ericaceous plant patches in the Arayu-Jigoku volcanic region, showing *Eubotryoides grayana* and *Rhododendron multiflorum* in a solfatara-field habitat and *Rhododendron multiflorum* in a forest-edge habitat. (B) Plant species sampled in this study. Four Ericaceae species, one Betulaceae species, and one Pinaceae species were collected across solfatara-field and forest-edge habitats at the two study sites, Arayu-Jigoku and Ofukasawa. The number of samples obtained for each species is shown in the bar plot.

At each study site, we established two belt zones (transects) representing contrasting habitats: (1) solfatara-field habitats, consisting of isolated plant patches on slopes near sulfur fumaroles, and (2) forest-edge habitats, consisting of patches located at least 3 m away from solfatara-field patches, adjacent to or within forested areas, and extending continuously for at least 3 m (Fig. 1A). In Arayu-Jigoku, we established 37 and 40 sampling positions in the solfatara-field and forest-edge habitats, respectively. In Ofukasawa, we established 15 sampling positions in each habitat. Sampling positions were separated by at least 3 m.

At each plant patch, approximately 2 cm³ of bulk soil was collected from a depth of 0–15 cm to measure soil pH in the field. Soil pH was measured using a portable pH meter, LAQUAtwin (HORIBA, Ltd., Japan), after moistening the soil with a small amount of water. In addition, bulk soil samples were collected from a depth of 0–15 cm at 28 representative sampling positions randomly selected across study sites and habitats, with at least five positions per site–habitat combination. These samples were used for soil chemical analyses and DNA metabarcoding after removing surface organic layers.

Root samples were collected from 304 individuals belonging to 15 plant species, mainly including five Ericaceae species and two ectomycorrhizal plant species in Betulaceae and Pinaceae that commonly occurred across the study sites and habitats. For each plant individual, ten root tips, each approximately 1 cm long, were collected. Sampling positions were recorded using the geographic positioning smartphone application Geographica (https://geographica.biz/): the altitude of sampling points ranged from 526 to 610 m. Root samples were immediately refrigerated in the field and stored at −50°C in the laboratory until DNA extraction.

To identify the plant species of root samples based on DNA sequences, leaves of the constituent plant species at the study sites were collected as reference material. Leaf samples were immediately refrigerated in the field and stored at −50°C in the laboratory.

### Measurements of soil chemical properties

Bulk soil samples were collected from the same depth as the target plant roots, 0–15 cm, at the 28 representative sampling points described above, after removing the surface organic layer. The samples were air-dried at room temperature for one week and then sieved through a 2-mm mesh.

Soluble cations were extracted from the soil as follows. First, 20 ml of 1 M CH₃COONH₄ (pH 7) was added to 1 g of air-dried soil, and the mixture was mechanically shaken at 80 rpm using a double shaker NR-30 (TIETECH, Japan) for 1 h. The suspensions were left to settle for 30 min, and the supernatants were filtered through glass filter paper (GA-55; ADVANTEC, Japan) using a disposable latex-free syringe (Terumo Syringe Lock Base; Terumo, Japan) and a filter holder (Swinex Filter Holder φ25 mm SX0002500; Merck Millipore Corporation, Germany).

Because trivalent aluminum, Al³⁺, is known to inhibit root elongation in plants growing in acidic soils (Horst et al., 2010), an additional extraction was performed to measure readily exchangeable Al³⁺ more accurately, following the protocol of a previous study [38]. Briefly, 30 ml of a mixed solution containing 1 M KCl and 0.5 M CaCl₂ was added to 3 g of air-dried soil, and the mixture was mechanically shaken at 80 rpm with a double shaker NR-30 (TIETECH, Japan) for 15 min. The suspensions were left to settle for 30 min, and the supernatants were filtered through a 0.45-μm cellulose acetate membrane filter (Minisart Syringe Filter S6555; Sartorius Stedim, Germany) using a latex-free syringe.

The concentrations of soluble base cations, Ca²⁺, Mg²⁺, K⁺, and Na⁺, and other soluble metal ions, Al³⁺, Mn²⁺, Co²⁺, Ni²⁺, Zn²⁺, and Pb²⁺, in the filtrates were measured using inductively coupled plasma optical emission spectrometry (iCAP PRO; Thermo Fisher Scientific, USA). The concentrations of soluble cations were treated as soil exchangeable cation contents. One sampling point was excluded from the analysis because of tube damage during transportation.

### Root and leaf DNA extraction and purification

Root samples for metabarcoding of the endophytic microbial community and leaf samples for plant species identification were processed separately for DNA extraction.

Surface soil of roots was removed by sonication in 1 mL of 0.5% Tween 20 at 45 kHz for 3 min. The roots were then surface sterilized by immersion in 3 mL of 1% (v/v) sodium hypochlorite solution (Nacalai Tesque, Japan; effective chlorine concentration approximately 11%) for 1 min, rinsed with 3 mL of sterile distilled water, and immersed in 99.5% ethanol for 30 s. The surface-sterilized roots were then stored at −20°C until further processing.

For root DNA extraction, the sterilized roots were freeze-dried overnight and pulverized at 6,500 rpm for 1 min in Lysing Matrix E tubes (MP Biomedicals, USA) using a Precellys Evolution Touch homogenizer (Bertin Technologies, France). The resulting powdered root samples were centrifuged at 3,010 × g for 3 min at 20°C, and the pellet was resuspended in 600 µl of cetyltrimethylammonium bromide buffer [100 mM Tris-HCl, pH 8.0; 1.4 M NaCl; 20 mM EDTA, pH 8.0; 2% cetyltrimethylammonium bromide]. The suspension was incubated at 60°C for 1 h and then centrifuged at 3,010 × g for 30 min at 20°C. After centrifugation, 200 µl of the supernatant was mixed with an equal volume of phenol/chloroform/isoamyl alcohol, 25:24:1, for 10 min and centrifuged at 3,010 × g for 30 min at 20°C. DNA in 80 µl of the resulting supernatant was precipitated by adding an equal volume of ice-cold 100% 2-propanol, followed by centrifugation at 17,800 × g for 10 min at 4°C. The DNA pellet was washed with ice-cold 70% ethanol and centrifuged again under the same conditions. After drying at 65°C for 30 min, the DNA was resuspended in 50 µl of 1× TE buffer [10 mM Tris-HCl, pH 8.0; 1 mM EDTA, pH 8.0] and used for downstream analyses.

Leaf samples were washed and pulverized using the same procedures as those applied to root samples. DNA was then extracted using a GENE PREP STAR PI-480α automated DNA extraction system with the Plant Tissue Reagent Kit ver. 1 (Kurabo Industries Ltd., Japan), according to the manufacturer’s protocol.

### Soil DNA extraction and purification

For DNA metabarcoding in soil, 20 soil samples were analyzed after excluding samples with insufficient material for DNA extraction to ensure reliable characterization of microbial communities [39]. The soil samples were thoroughly mixed, and plant debris and pebbles were removed from approximately 10–40 g of moist soil. The soil was then frozen at −50°C and freeze-dried overnight. After freeze-drying, 2 g of each dried soil sample was passed through a 0.5-mm mesh sieve, and fine roots were removed with tweezers.

The sieved soil was transferred to a 50-ml centrifuge tube containing 4-mm, 1-mm, and 0.5-mm zirconium beads (AS ONE, Japan). The tubes were shaken at 6.5 m s⁻¹ for 60 s using a FastPrep-24 instrument (MP Biomedicals, USA). After bead-beating, 6 ml of extraction buffer [10 mM Tris-HCl, pH 8.0; 1 mM EDTA; 1% Triton X-100] was added, and the mixture was vortexed for 5 s. The samples were then centrifuged at 20,000 × g for 10 min, and 500 µl of the supernatant was subjected to DNA extraction using the Extrap Soil DNA Kit Plus Ver. 2 (BioDynamics Laboratory Inc., Japan) without the bead-beating step included in the manufacturer’s protocol.

### PCR amplification

Amplicon libraries were prepared using a two-step PCR protocol. In the first PCR, locus-specific regions were amplified using primers containing partial Illumina adapter sequences and 1–6 random nucleotides to improve sequencing quality. In the second PCR, full-length Illumina adapter sequences and sample-specific 8-mer indices were added using fusion primers.

The primers used for the first PCR shared a common structure consisting of partial Illumina adapter sequences, followed by 1–6 random nucleotides and the locus-specific primer sequences. The forward primer structure was 5′-TCG TCG GCA GCG TCA GAT GTG TAT AAG AGA CAG - [1–6 random nucleotides] - [locus-specific sequence]-3′, and the reverse primer structure was 5′-GTC TCG TGG GCT CGG AGA TGT GTA TAA GAG ACA G - [1–6 random nucleotides] - [locus-specific sequence]-3′.

Prokaryotic communities in root and soil samples were characterized by amplifying the V5– V6 region of the prokaryotic 16S rRNA gene using the primers 799F [40] and 1107R [41]. PCR reactions were performed using the KOD One system (Toyobo, Japan), with each primer at a final concentration of 0.3 µM. The thermal cycling conditions were 35 cycles of 98°C for 10 s, 60°C for 5 s, and 68°C for 5 s, followed by a final extension at 68°C for 2 min. The ramp rate was set to 1°C s⁻¹ to reduce chimera formation [42].

Fungal communities were analyzed by amplifying the ITS1 region using the primers ITS1F-KYO1 and ITS2-KYO2 [43]. PCR reactions were performed using the Ampdirect Plus PCR Kit (Shimadzu, Japan), with each primer at a final concentration of 0.3 µM. The thermal cycling conditions were 32 cycles of 94°C for 30 s, 52°C for 1 min, and 72°C for 1 min, followed by a final extension at 72°C for 7 min. The ramp rate was also set to 1°C s⁻¹.

Plant species in root and reference leaf samples were analyzed by amplifying the ITS2 region using the primers ITS-3p62plF1 and ITS-4unR1 [44], under the same PCR conditions as those used for fungal ITS amplification.

For the second PCR, fusion primers consisted of a P5 or P7 adapter sequence, an 8-mer sample index, and partial sequencing primer sequences [45]. The forward fusion primer consisted of the P5 adapter, an 8-mer index, and a partial sequencing primer sequence, 5′-AAT GAT ACG GCG ACC ACC GAG ATC TAC AC - [8-mer index] - TCG TCG GCA GCG TC-3′. The reverse fusion primer consisted of the P7 adapter, an 8-mer index, and a partial sequencing primer sequence, 5′-CAA GCA GAA GAC GGC ATA CGA GAT - [8-mer index] - GTC TCG TGG GCT CGG-3′. Secondary PCR was performed using KOD One PCR Master Mix under the following conditions: 15 cycles of 98°C for 10 s, 55°C for 5 s, and 68°C for 5 s, followed by a final extension at 68°C for 2 min. The ramp rate was set to 1°C s⁻¹.

PCR amplicons were purified and normalized using AMPure XP beads (Beckman Coulter, USA) at a 3:5 reagent-to-DNA ratio to remove fragments shorter than 200 bp. The amplicons were then size-selected using E-Gel SizeSelect 2 (Invitrogen, USA), targeting approximately 480 bp for prokaryotic and fungal amplicons and approximately 520 bp for plant amplicons. Sequencing libraries for the prokaryotic 16S rRNA gene, fungal ITS1 region, and plant ITS2 region were processed on an Illumina MiSeq platform with a 20% PhiX spike-in. To maximize sequence quality, only forward reads were generated using 301 forward cycles.

### Bioinformatics

#### Sequence processing and microbial taxonomic assignment

Sequence data were processed following previously described metabarcoding workflows [46–48]. Prokaryotic 16S rRNA gene and fungal ITS1 sequences were analyzed to characterize microbial communities, whereas plant ITS2 sequences were used exclusively to identify the host plant of each root sample. Briefly, raw data were converted into FASTQ files using bcl2fastq v1.8.4 and demultiplexed using Claident v0.9.2022.01.26 [49, 50]. Following target-sequence extraction with Cutadapt v4.4 [51], reads were filtered and trimmed using DADA2 v1.18.0 in R v4.3.1 (maxN = 0, maxEE = 2, minLen = 100, minQ = 10, truncQ = 11, truncLen = 0, and rm.phix = TRUE) [52, 53]. Amplicon sequence variants (ASVs) were inferred within sequencing runs, and the resulting sequence tables were combined using mergeSequenceTables function in DADA2 [52, 53]. Chimeric sequences were subsequently removed using “removeBimeraDenovo” function in DADA2 with the consensus method [52, 53]. After chimera removal, the datasets contained 4,078,351 prokaryotic reads and 6,865,366 fungal reads.

Potential contaminants in the prokaryotic 16S rRNA gene and fungal ITS1 datasets were identified using the prevalence-based “isContaminant” function in the R package decontam v 1.30.0 [54]. Decontamination was not performed for plant ITS2 data because enough negative controls were not detected to estimate contamination.

Prokaryotic and fungal ASVs were taxonomically assigned using “assignTaxonomy” function in DADA2 against SILVA v138.2 for prokaryotes [55] and the UNITE General FASTA release version 10.0 for fungi [56], respectively. ASVs were then clustered into operational taxonomic units (OTUs) at 97% sequence identity using VSEARCH v2.15.2 [57], and each OTU inherited the taxonomic assignment of its representative ASV. Chloroplast and mitochondrial OTUs were excluded from the prokaryotic dataset, and only OTUs assigned to Fungi were retained in the fungal dataset.

#### Host plant identification

Host plant identity was determined for each root sample using plant ITS2 sequences and a reference leaf ITS2 library. Because multiple ITS2 OTUs were sometimes recovered from a single reference leaf sample owing to sequence length and intragenomic variation [58], root-derived plant OTUs were compared with all reference leaf OTUs using local BLAST searches implemented in NCBI BLAST+ v2.17.0+ [59]. Matches with ≥ 97% sequence identity over an alignment length of ≥ 100 bp were used to identify candidate host taxa.

To classify plant OTUs unresolved by the reference leaf library and evaluate their consistency with host candidates identified from the leaf library, secondary taxonomy was obtained from assignments to their representative ASVs using Claident v0.9.2022.01.26 [49] and its “overall_genus” database. Assignments were generated using the query-centric auto-k-nearest-neighbor (QCauto) method under strict and relaxed criteria and a nearest-neighbor method [50] with the setting “5,90%”, then integrated in that order of priority. Additional manual BLAST searches (https://blast.ncbi.nlm.nih.gov/) against the NCBI nucleotide database were conducted on June 6 and July 28, 2026 [60]. These secondary assignments were not used independently to establish host identity.

To minimize potential contamination and retain samples with a clear dominant host signal, only samples in which a single taxonomic group accounted for ≥ 90% of total plant ITS2 reads and its identity was supported by the reference leaf library were retained. These samples were assigned the corresponding host identity based on the reference leaf library. For this filtering, reads were summed across OTUs assigned to each plant genus, except where noted below, based on both leaf-library and secondary assignments, and their proportions of total plant ITS2 reads were calculated. Reads unresolved at the applicable taxonomic level were included in the denominator.

Since some taxa could not be distinguished using ITS2 sequences, *Rhododendron multiflorum* and *R. japonicum* were treated collectively as *Rhododendron* spp. Similarly, *Eubotryoides grayana* and *Gaultheria adenothrix* shared indistinguishable ITS2 sequences; reads assigned to these genera or only to their tribe Gaultherieae were therefore pooled at the tribe level when applying the 90% criterion. Samples meeting this criterion were labeled as *E. grayana*, because vegetation surveys showed that *E. grayana* overwhelmingly dominated over *G. adenothrix* in the study areas. Host categories represented by fewer than 10 samples were excluded, leaving six focal taxa: *Enkianthus campanulatus*, *Rhododendron* spp., *Eubotryoides grayana*, *Vaccinium smallii*, *Betula ermanii*, and *Pinus parviflora* (Fig. 1B).

#### Preparation of datasets for downstream analyses

To ensure sufficient sequencing depth before coverage-based rarefaction, root samples with fewer than 2,000 reads and soil samples with fewer than 5,000 reads were excluded before rarefaction based on OTU rarefaction curves generated using “rarecurve” function (Fig. S1). For each prokaryotic or fungal root or soil dataset, curve slopes were calculated using “rareslope” in vegan v2.7-2 [61]. To account for differences in the sequencing depth at which OTU diversity saturates, the largest slope near the samples’ full observed sequencing depths was used as a common slope threshold, representing the least flattened curve. Sample-specific depths were selected where slopes reached or fell below this threshold, and random subsampling was performed using “rrarefy” with a seed. Dataset preparation and all downstream statistical analyses were conducted in R v4.5.3 [53].

The root-associated prokaryotic and fungal datasets each included 206 samples from 105 sampling positions, with 205 samples shared between the two datasets. Both prokaryotic and fungal soil datasets included the same 20 representative samples. Following rarefaction and sample selection, the root-associated datasets contained 612 prokaryotic OTUs (585 bacterial, 1 archaeal, and 26 unclassified) and 484 fungal OTUs. The soil datasets contained 490 prokaryotic OTUs (487 bacterial, 2 archaeal, and 1 unclassified) and 471 fungal OTUs.

All OTUs were retained for alpha diversity analyses, whereas OTUs detected in fewer than three samples were excluded from community structure and habitat/host preference analyses, given the limited information available for preference estimation from such infrequent occurrences and to maintain consistent filtering criteria across these analyses (Fig. S2). Samples retaining no OTUs after filtering were excluded from the corresponding analyses. Sensitivity to the prevalence threshold was assessed by comparing preference scores for shared OTUs with those obtained using alternative thresholds of one and five samples (Figs. S3–4).

#### Statistical analysis for soil chemical profiling

To characterize patterns of soil chemical variation across habitats, principal component analysis (PCA), permutational multivariate analysis of variance (PERMANOVA) [62], and permutational analysis of multivariate dispersions (PERMDISP) [63] were performed using soil pH and exchangeable cation concentrations. PERMANOVA was performed based on Euclidean distances of continuous environmental variables using the “adonis2” function in the R package vegan v2.7-2 [64], with 9,999 permutations. The effect of habitat was evaluated while incorporating study site as a stratification factor. Homogeneity of multivariate dispersion among habitats was assessed using PERMDISP with the “betadisper” function in vegan. In these multivariate analyses, each environmental variable was standardized (mean = 0, standard deviation = 1) to minimize bias arising from differences in measurement scales.

For each chemical parameter, Wilcoxon rank-sum tests were conducted to evaluate differences between solfatara-field and forest-edge habitats within each study site using the “wilcox_test” function in the R package rstatix v 0.7.3 [65]. The resulting *P*-values were adjusted using the Benjamini-Hochberg correction [66] to control the false discovery rate (FDR).

Prior to statistical analyses, soil elemental concentrations were blank corrected by subtracting the mean concentration of blank samples for each element. Co²⁺ and Ni²⁺ were excluded from subsequent analyses because more than half of the measurements were below the detection limits. Three of the 28 sampling points were excluded from the multivariate analyses because of tube breakage for Al³⁺ measurement (*n* = 1) or non-detect measurements for Pb²⁺ (*n* = 2). For the univariate analyses, all available measurements from the 28 sampling points were retained, with missing values resulting only from non-detect measurements.

#### Community differentiation

To identify the factors shaping root-associated microbial community differentiation, we evaluated the effects of habitat, host plant identity, and sample type (root or soil) on microbial diversity and community composition.

Alpha diversity indices, namely prokaryotic and fungal OTU richness, were compared between solfatara-field and forest-edge habitats within each study site using Wilcoxon rank-sum tests implemented with the “wilcox_test” function in the rstatix package [65]. The resulting *P*-values were adjusted using the Benjamini-Hochberg correction. Differences in OTU richness between root and soil samples were evaluated using a negative binomial generalized linear mixed-effects model (GLMM), implemented with the R package glmmTMB v 1.1.14 [67], with sample type as a fixed effect and sampling position as a random effect to account for the non-independence of samples collected from the same position. A negative binomial distribution with a linear mean– variance relationship (nbinom1) was selected based on model diagnostics using the R package DHARMa v 0.5.0 [68].

To assess how microbial community composition varied with habitat, host plant identity, and sample type, we analyzed two datasets separately: (i) root-associated microbial communities from all sampling points, focusing on habitat and host identity, and (ii) 20 representative soil samples together with root samples collected at those sampling positions, focusing on habitat and sample type. The root–soil dataset comprised soil samples from 20 positions and root samples from 19 of those positions, retaining all soil samples despite missing root pairs. For each dataset, community structure was visualized using principal coordinate analysis (PCoA) based on Sørensen dissimilarities calculated from presence–absence community matrices. Differences in community composition were assessed using PERMANOVA [62], and homogeneity of multivariate dispersions was evaluated using PERMDISP [63]. Variation partitioning was also performed to quantify the unique and shared contributions of the explanatory variables using the “varpart” function in the R package vegan v2.7-2 [64].

For the root-only dataset, PERMANOVA was used to assess the effects of habitat, host plant identity, and their interaction. Sequential tests were performed with habitat entered before host plant identity, reflecting the broader ecological scale of habitat-level variation. For the root–soil dataset, habitat effects were tested separately within each sample type, whereas root–soil differences were tested separately within each habitat.

To account for the spatial structure of the sampling design, permutations in PERMANOVA and PERMDISP were restricted within study sites for comparisons of habitats and host plant identities, and within sampling positions for root–soil comparisons. All tests were based on 9,999 permutations. For all PERMDISP analyses, distances to group spatial medians were calculated using “betadisper” function with bias.adjust = TRUE in the R package vegan [64]. Within PERMANOVA and PERMDISP separately, *P* values for the two habitat comparisons were adjusted together, and those for the two root–soil comparisons were adjusted together, using the Benjamini– Hochberg FDR method [66].

#### OTU-level habitat and host specialization

To identify the OTU-level niche specialization underlying the observed habitat- and host-associated community differentiation, we quantified habitat and host niche preferences for individual root-associated prokaryotic and fungal OTUs using the following randomization-based analyses. Habitat and host preferences were evaluated independently by controlling for the other factors during randomization, allowing us to distinguish OTUs associated with a particular habitat, host identity, or both. After calculation of these preference indices, we evaluated the association between habitat and host specialization for prokaryotes and fungi separately, using Spearman’s rank correlation with “cor.test” function in the R package stats v4.5.3, excluding OTUs with missing values for either index. Habitat specialization was represented by the absolute *z*-standardized 2DP score, reflecting the strength of preference for either the solfatara field or the forest edge. Host specialization was represented by the signed *z*-standardized *d′* index, with higher values indicating greater specialization relative to the null expectation.

#### Habitat preference analysis

Habitat preference was quantified using the two-dimensional preference (2DP) index [35]. Presence–absence community matrices were used to calculate a *z*-standardized habitat association score for each microbial OTU by comparing the observed occurrence frequency in each habitat with a null distribution generated by randomizing habitat labels while preserving host plant identity and study site. For each habitat *i* and microbial OTU *j*, a *z*-standardized habitat association score was calculated as:

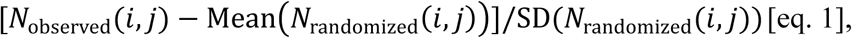

where *N*_observed_(*i*, *j*) represents the number of root samples in which habitat *i* and OTU *j* co-occurred in the original dataset, and Mean(*N*_randomized_(*i*, *j*)) and SD(*N*_randomized_(*i*, *j*)) are the mean and standard deviation of co-occurrence counts across randomized matrices. Statistical significance was evaluated using two-tailed tests with 99,999 permutations, followed by Benjamini-Hochberg correction [66].

#### Host preference analysis

To assess host preferences of each microbial OTU and microbial preferences of each host, host– microbe association specificity was quantified in both directions using the *d′* index of association specificity with the “dfun” function [69] in the R package bipartite v2.23 [70].

For microbial OTUs, host preference was quantified by calculating *d′* for each prokaryotic and fungal OTU. To assess statistical significance, we randomly reassigned host plant labels among root samples collected from the same study site and habitat for the presence–absence community matrix. Observed *d’* values were compared with a null distribution generated from 99,999 randomizations. For each prokaryotic and fungal OTU, a *z*-standardized host preference score was calculated as:

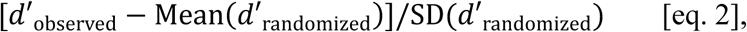

where *d*′_observed_ is the *d′* value calculated from the original data matrix, and Mean(*d*′_randomized_) and SD(*d*′_randomized_) are the mean and standard deviation of *d′* values obtained from randomized matrices. Statistical significance was assessed using two-tailed tests followed by Benjamini-Hochberg correction. Conversely, microbial preference was quantified by calculating *d′* for each host plant identity, using the same procedures as above. Statistical significance was evaluated using 9,999 permutations to balance computational cost with the resolution required for multiple-testing correction.

Furthermore, to identify which host–microbial associations underlie the patterns observed in the specificity analysis, pairwise associations between individual host plant identities and microbial OTUs were evaluated using the *z*-standardized 2DP index [eq. 1]. Statistical significance was assessed using two-tailed tests based on 99,999 permutations, followed by Benjamini-Hochberg correction.

#### Phylogenetic signals in habitat and host preferences

To determine whether habitat and host specialization of root-associated fungi reflects phylogenetic constraints, we evaluated phylogenetic signals in preference traits among closely related fungal taxa. We focused on the family Hyaloscyphaceae (Helotiales), whose OTUs exhibited substantial variation in habitat and host preferences, enabling us to test whether niche specialization was conserved among related lineages.

Phylogenetic relationships of Hyaloscyphaceae OTUs were reconstructed using 24 OTU sequences identified with the UNITE fungal database version 10.0 [56] together with 14 reference sequences retrieved from the NCBI database. Sequences were aligned using MAFFT v7.526 [71], and poorly aligned regions were removed using trimAl v1.5.1 with a gap threshold of 0.6 [72]. A Neighbor-Joining phylogenetic tree was constructed in MEGA12.1 [73] using the Kimura 2-parameter model [74], with gaps and missing data treated by pairwise deletion. The tree was rooted using *Hymenoscyphus fraxineus* (GenBank accession no. PX574263) as the outgroup. Branch support was evaluated using 1,000 bootstrap replicates. The resulting tree was visualized with habitat preference scores, host preference scores, and their significance levels mapped onto the tips using the R packages ape v5.8. [75] and ggtree v3.14.0 [76].

Phylogenetic signals were evaluated for Hyaloscyphaceae OTUs detected in this study, for which habitat and host preference scores were available. Phylogenetic signals in preference traits were respectively evaluated using Blomberg’s *K* [77] and Pagel’s *λ* [78] implemented in the “phylosig” function of the R package phytools v2.5.2 [79]. Statistical significance of Blomberg’s *K* was assessed using 9,999 randomizations, whereas that of Pagel’s *λ* was assessed using a likelihood-ratio test. Blomberg’s *K* quantifies the similarity of trait values among closely related taxa relative to expectations under Brownian-motion trait evolution; values near 1 indicate Brownian-motion evolution, values near 0 indicate phylogenetic independence, and values greater than 1 indicate stronger phylogenetic conservatism than expected under Brownian motion. Pagel’s *λ* estimates the extent to which phylogenetic covariance explains trait variation, with values near 0 indicating phylogenetic independence and values near 1 indicating a Brownian-motion-like covariance structure. Analyses were performed separately for habitat preference (2DP score for the solfatara habitat) and host preference (*z*-standardized *d′* score).

## Results

### Soil chemical characterization in solfatara fields

Soil chemical properties differed between solfatara-field and forest-edge habitats across the two study sites, although the magnitude of these differences varied among sites and elements. In the PCA, the first two principal components explained 53.2% and 15.3% of the total variance, respectively (Fig. 2A). Loading vectors indicated that PC1 was primarily associated with exchangeable cation concentrations, including K⁺, Mg^2^⁺, Al^3^⁺, and Na⁺, whereas PC2 mainly reflected variation in Mn²⁺ and Ca²⁺, with a strong positive loading, and soil pH and Pb^2+^, with a negative loading (Table S1).

**Fig 2.**
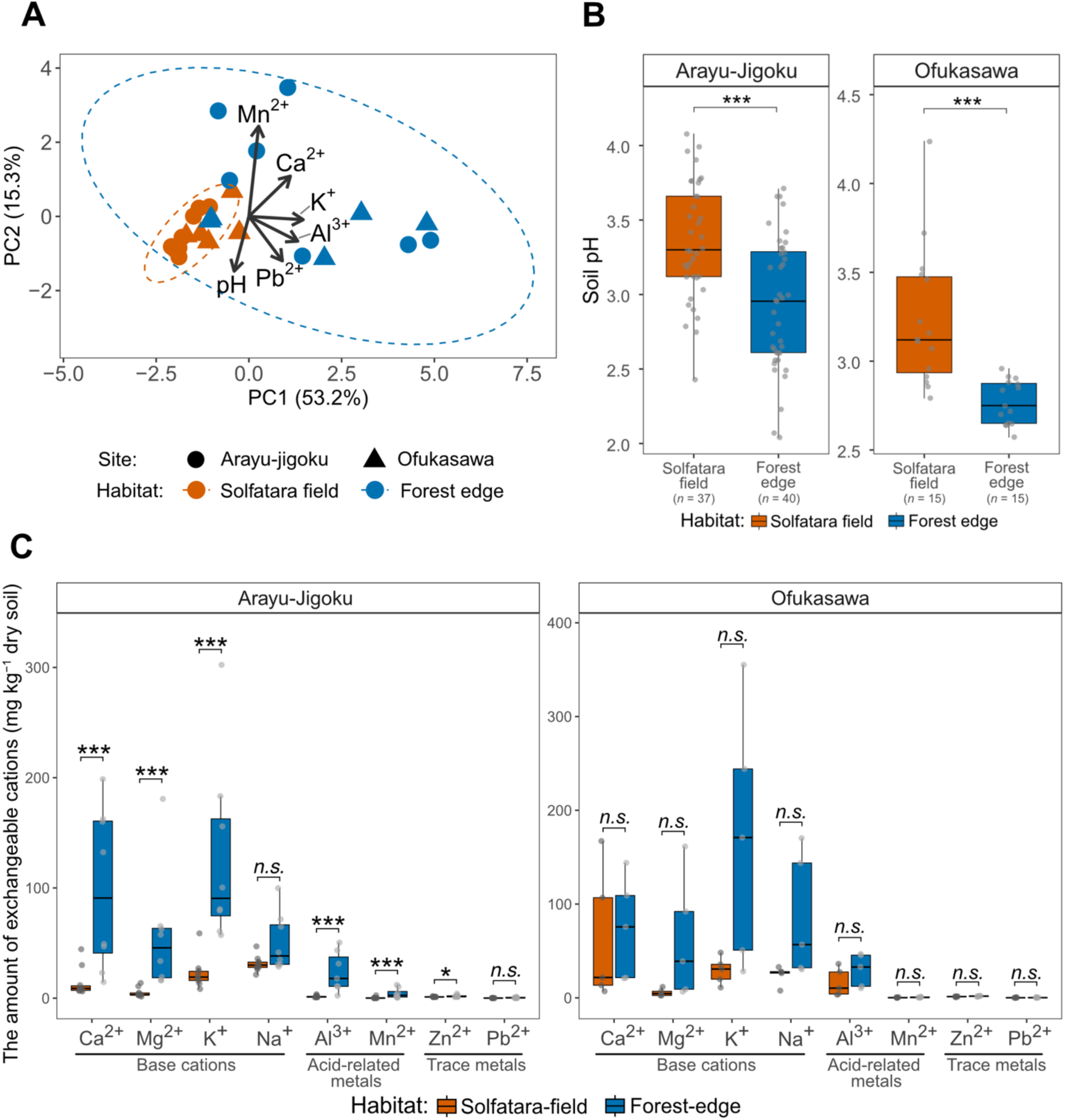
Comparison of soil chemical properties between solfatara-field and forest-edge habitats. (A) Multivariate analysis of soil cation concentrations. The pH and cation concentrations of soil samples collected from 28 representative sampling positions are shown based on a principal component analysis (PCA). Arrows indicate the direction and magnitude of the loadings of each cation on the first two principal components. The dashed lines represent the 95% confidence intervals of solfatara-field and forest-edge samples. (B) Comparison of pH between solfatara-field and forest-edge habitats at each study site. The sample size for each combination is indicated in parentheses. (C) Comparison of cation concentrations (Ca^2+^, Mg^2+^, K^+^, Na^+^, Al^3+^, Mn^2+^, Zn^2+^, Pb^2+^) between solfatara-field and forest-edge habitats at each study site (*n* = 10 in the solfatara field and *n* = 8 in the forest edge at Arayu-jigoku for all cations except Al³⁺, for which *n* = 10 and *n* = 7, respectively; *n* = 5 in each habitat at Ofukasawa).

Consistent with these ordination patterns, PERMANOVA detected significant differences in soil chemical composition between habitats (Table S2; *R²* = 0.28, *P* < 0.001). PERMDISP also revealed significant differences in within-group dispersion among habitats (Table S3; *P* < 0.001), indicating that the multivariate separation was partly associated with differences in heterogeneity.

At the level of individual variables, soil pH was significantly higher in solfatara-field than in forest-edge habitats at both study sites, although soils in both habitats remained strongly acidic, with values around pH 3.0 (rank-biserial correlation, *r_rb_* = 0.43 at Arayu-jigoku and 0.72 at Ofukasawa; Wilcoxon rank-sum test, both FDR < 0.001; Fig. 2B). At Arayu-Jigoku, concentrations of multiple cations, including major base cations (Ca²⁺, Mg²⁺, and K⁺) and metal cations that can be more soluble under acidic soil conditions (Al³⁺ and Mn²⁺, and Zn²⁺), were significantly lower in solfatara-field than in forest-edge habitats (*r_rb_* = 0.59–0.84; Wilcoxon rank-sum test, all FDR < 0.05; Fig. 2C). Although no individual element showed a statistically significant difference between the habitats at Ofukasawa, most variables displayed directional trends consistent with those observed at Arayu-Jigoku.

Collectively, these results indicate that, despite strong acidity in both habitats, solfatara-field soils were characterized by relatively high pH and severe cation depletion, including reduced concentrations of both base cations and aluminum. In contrast, forest-edge soils were more acidic but retained higher concentrations of nutrients and metals, while also showing greater within-habitat heterogeneity. These patterns suggest that the two habitats differed not simply in stress intensity, but also in the types of soil chemical constraints imposed on plant and microbial communities.

### Community composition and diversity patterns

Prokaryotic community composition differed between root and soil samples and between solfatara-field and forest-edge habitats across the two study sites (Fig. 3A; Supplementary Fig. S5A). Root-associated communities were consistently dominated by actinobacterial lineages, particularly Frankiales, which accounted for approximately 50% of total reads across sites and habitats. At both study sites, roots in solfatara-field habitats showed higher relative abundances of Catenulisporales (18.3% at Arayu-Jigoku and 8.8% at Ofukasawa), whereas forest-edge roots showed higher relative abundances of Mycobacteriales (12.5% and 25.0%, respectively) (Fig. 3A). Soil communities also differed between habitats: Frankiales (25.3% and 29.1%) together with Acidimicrobiales (17.6% and 15.3%) predominated in solfatara-field soils, whereas Mycobacteriales was the dominant order in forest-edge soils (23.6% and 58.1%, respectively; Fig. S5A).

**Fig 3.**
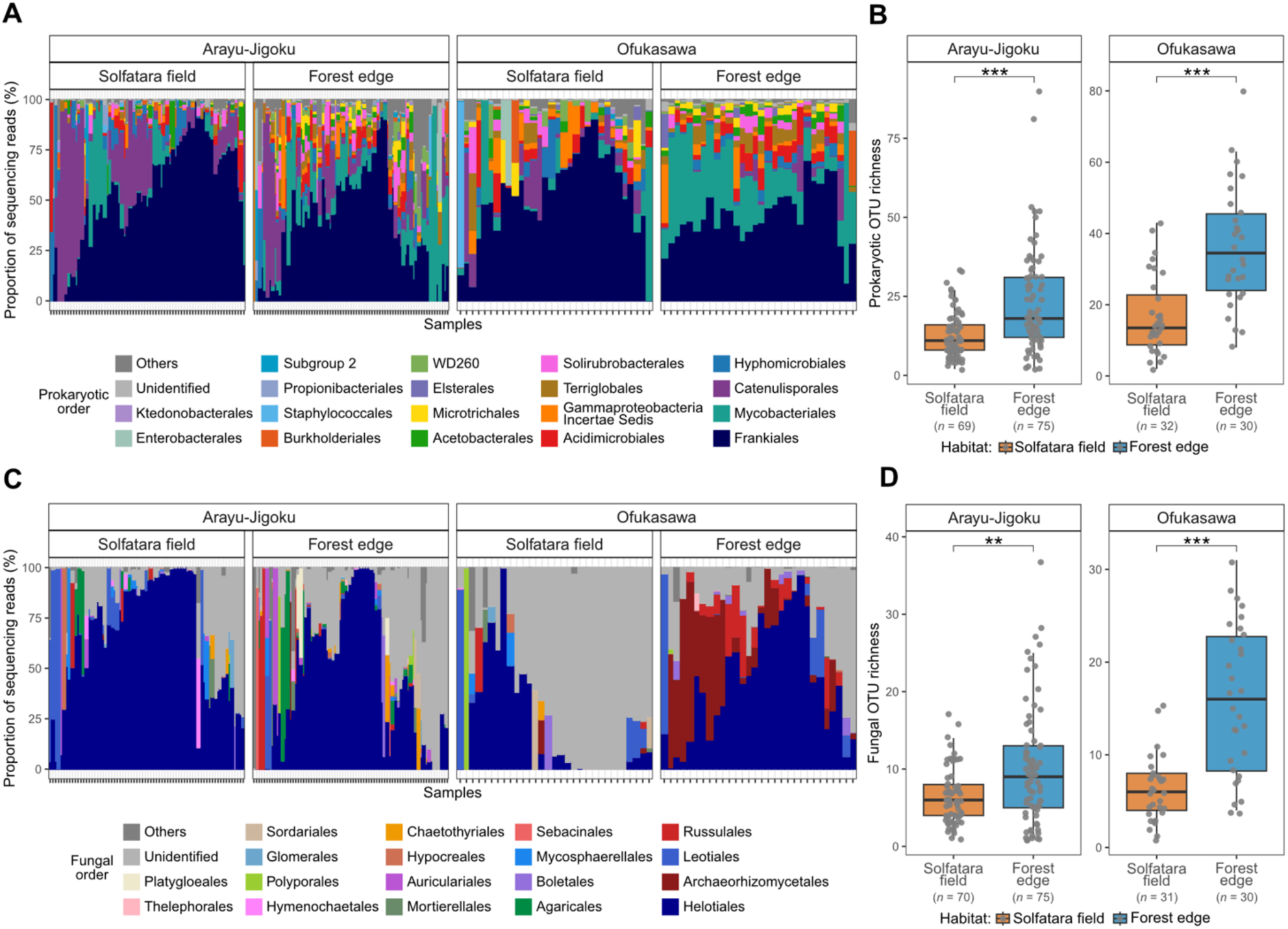
Root-associated prokaryotic and fungal community compositions and OTU richness. (A, C) Order-level taxonomic composition of prokaryotic (A) and fungal (C) communities. Root samples were ordered depending on a clustering approach based on Bray-Curtis dissimilarity between samples. (B, D) Comparison of OTU richness of prokaryotes (B) and fungi (D) between solfatara-field and forest-edge habitats.

Comparison of root samples with corresponding soil samples showed that root-associated communities reflected environmental differences observed in soil, while also being characterized by increased relative abundances of specific taxa, particularly Frankiales, resulting in community composition distinct from that of corresponding soil samples (Fig. S5A). This root–soil differentiation was accompanied by reduced OTU richness in roots relative to soils. Across corresponding sampling positions, prokaryotic OTU richness was significantly higher in soil than in root samples, with soil samples exhibiting 3.70-fold greater richness (95% CI: 2.89–4.73; negative binomial GLMM, *P* < 0.001; Fig. S5B). Within root samples, OTU richness was significantly lower in solfatara-field habitats than in forest-edge habitats (*r_rb_* = −0.44 and −0.67; Wilcoxon rank-sum test, FDR < 0.001; Fig. 3B), whereas soil OTU richness did not differ significantly between habitats (*r_rb_* = 0.20 and 0.88; Wilcoxon rank-sum test, FDR > 0.05).

Fungal community composition differed between solfatara-field and forest-edge habitats in both root and soil samples, although the dominant taxa and the magnitude of these differences varied between study sites (Fig. 3C; Fig. S5C). In roots, fungal communities were generally dominated by Helotiales. At Arayu-Jigoku, forest-edge roots showed higher relative abundances of basidiomycete lineages, including Agaricales (7.3%) and Boletales (4.5%), than solfatara-field roots (Fig. 3). At Ofukasawa, solfatara-field roots contained a high proportion of unidentified ascomycete taxa, whereas forest-edge roots showed marked increases in Helotiales (48.5%), Archaeorhizomycetales (21.2%), and Russulales (8.0%). Soil fungal communities contained a broader range of abundant fungal lineages than root-associated communities (Fig. S5C). At Arayu-Jigoku, soil communities were characterized by ascomycete-dominated assemblages in solfatara-field habitats and by increased relative abundances of basidiomycete orders, particularly Agaricales (18.8%) and Archaeorhizomycetales (12.5%), in forest-edge habitats. At Ofukasawa, soil communities showed a comparable habitat-dependent pattern, with forest-edge soils being strongly dominated by Helotiales (62.2%) together with Archaeorhizomycetales (13.5%).

Comparison of root samples with corresponding soil samples likewise showed that root-associated fungal communities reflected environmental differences in the surrounding soil, while also being characterized by increased relative abundances of specific taxa, especially Helotiales (Fig. S5C). This root–soil differentiation was accompanied by reduced fungal OTU richness in roots relative to soil samples. At corresponding sampling points, fungal OTU richness was substantially higher in soil than in root samples, with soil richness estimated to be approximately fivefold higher than root richness (Soil/Root ratio = 4.96, 95% CI: 3.92–6.29; negative binomial GLMM, *P* < 0.001; Fig. S5D). Within root samples, fungal OTU richness was significantly lower in solfatara-field habitats than in forest-edge habitats (*r_rb_* = −0.28 and −0.71; Wilcoxon rank-sum test, FDR < 0.001; Fig. 3D), whereas soil fungal communities showed the same tendency but did not differ significantly between habitats (*r_rb_* = −0.84 and −0.68; Wilcoxon rank-sum test, FDR > 0.05).

### Community structural differentiation across habitats, hosts, and sample types

PCoA and PERMANOVA showed that both prokaryotic and fungal root-associated communities were significantly structured by habitat and host identity (Fig. 4A–B; Table S4). Habitat (prokaryotes: *R²* = 0.052; fungi: *R²* = 0.066; both *P* < 0.001) and host identity (prokaryotes: *R²* = 0.066; fungi: *R²* = 0.083; both *P* < 0.001) explained comparable proportions of community variation, although host identity consistently accounted for slightly more variation even when habitat was fitted first in the sequential PERMANOVA. A significant interaction between habitat and host identity was detected only for fungal communities (*R²* = 0.029, *P* < 0.05), but its contribution was small relative to the single effects (Table S4). Variation partitioning showed that habitat and host identity explained largely independent fractions of variation, with negligible overlap (Table S5). PERMDISP detected no differences in dispersion among habitats (both *P* > 0.05; Tables S6) but significant heterogeneity among host plants (both *P* < 0.01; Tables S6–7).

**Fig 4.**
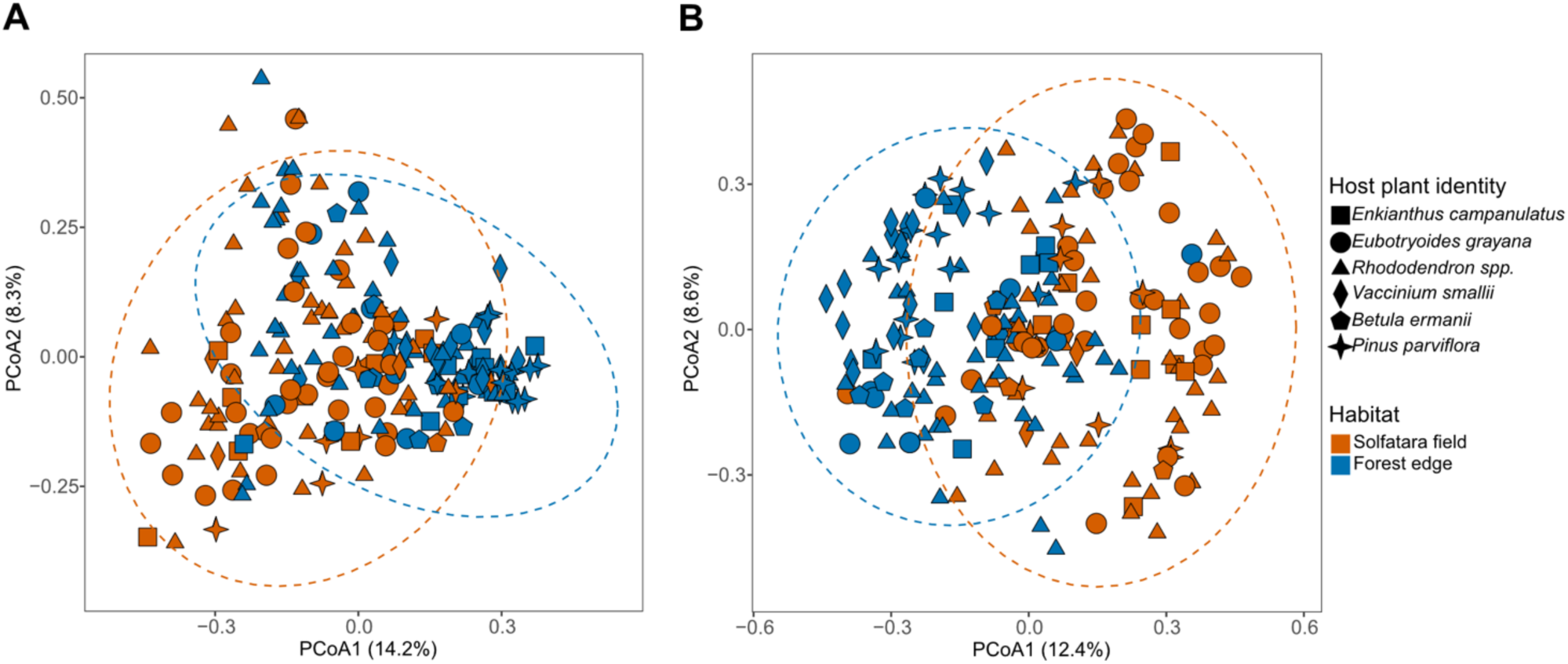
Variation in root-associated prokaryotic and fungal community structure across habitats and host plant identities. Principal coordinate analysis (PCoA) plots based on the Sørensen dissimilarity index show community compositional variation in root samples for (A) prokaryotes and (B) fungi. Colors represent habitats and symbols indicate host plant identity. Dashed ellipses indicate 95% confidence intervals for each habitat: red, solfatara field; blue, forest edge.

Focusing on the representative root and soil samples, we evaluated habitat effects separately for root and soil communities, as well as root–soil differentiation within each habitat. Prokaryotic and fungal community compositions differed significantly between habitats in both sample types (PERMANOVA, *R²* = 0.081–0.291, all FDR < 0.001; Fig. S6; Table S8), and distinct root–soil differentiation was observed within both habitats (PERMANOVA, *R²* = 0.106–0.279, all FDR < 0.001; Fig. S6; Table S8). Variation partitioning consistently revealed that habitat and sample type explained largely independent fractions of community variation across both microbial groups (Table S9). Furthermore, PERMDISP analysis indicated no significant differences in dispersion between habitats for both root and soil subsets (both FDR > 0.05; Fig. S6; Tables S10), whereas dispersion differed significantly between sample types (root–soil) within each habitat for both prokaryotic and fungal communities (both FDR < 0.001; Fig. S6; Tables S10).

### OTU-level preferences for habitats and host plants

Within the root-associated prokaryote community, 23 OTUs showed significant habitat preferences, but none showed significant preference for solfatara-field habitats (Fig. 5A). Instead, these OTUs were biased toward forest-edge habitats, as exemplified by Acidobacteriaceae [P_0205]. Host preference was detected for 9 prokaryotic OTUs, including *Acidothermus* [P_0093] and *Candidatus* Liberibacter [P_3583]. Only two prokaryotic OTUs, *Acidothermus* [P_0182] and *Mycobacterium* [P_0230], showed significant preferences in both analyses, being associated with forest-edge habitats and particular host plant identities.

**Fig 5.**
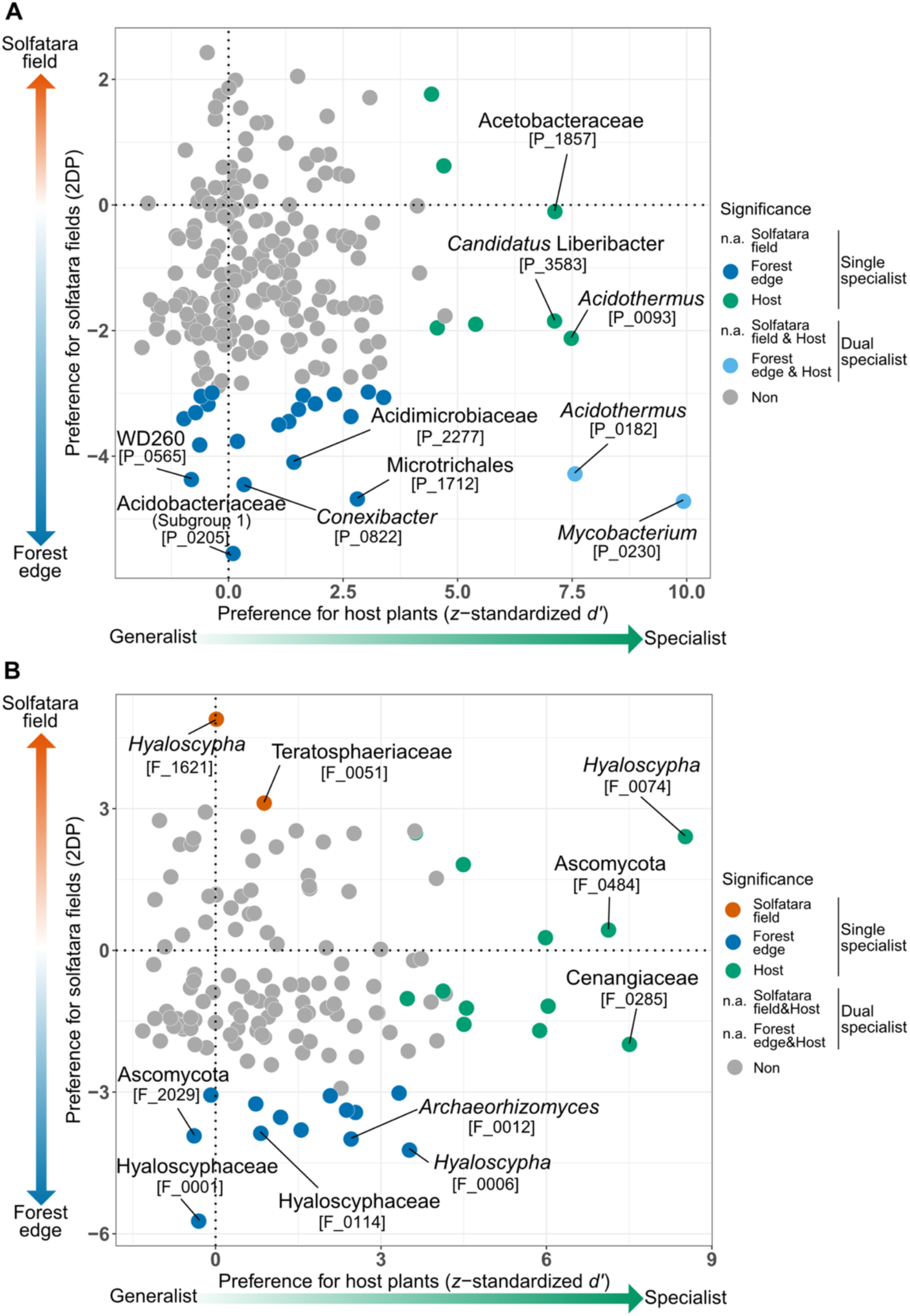
Host preference and solfatara-field preference of prokaryotic and fungal OTUs. Host preference, measured as *z*-standardized *d′*, is shown on the vertical axis, and solfatara-field preference, measured as the *z*-standardized two-dimensional preference index (2DP), is shown on the horizontal axis for (A) prokaryotic and (B) fungal OTUs. Statistical significance was assessed using two-tailed tests followed by false discovery rate (FDR) correction with the Benjamini–Hochberg method. Colors indicate significant associations: blue, host plant identity; red, habitat; yellow, both host plant identity and habitat; gray, not significant.

Within the root-associated fungal community, 2 OTUs, namely, *Hyaloscypha* [F_1621] and Teratosphaeriaceae [F_0051], exhibited significant preference for solfatara-field habitats, whereas 13 OTUs, including five Hyaloscyphaceae fungi [F_0001, F_0006, F_0019, F_0114, F_0271], significantly preferred forest-edge habitats (Fig. 5B). Host preference was detected for 12 fungal OTUs, including *Hyaloscypha* [F_0074] and Cenangiaceae [F_0285]. However, no fungal OTU showed statistically significant patterns in both habitat-preference and host-preference analyses, indicating that fungal OTUs associated with particular habitats were distinct from those associated with particular host plant identities. Notably, several Hyaloscyphaceae OTUs displayed contrasting habitat preferences despite low host specificity, suggesting niche differentiation within this family.

In each dataset of prokaryotes and fungi, habitat specialization, measured as the absolute *z*-standardized 2DP score, and host specialization, measured as *z*-standardized *d′*, were not significantly correlated in either prokaryotes or fungi (Spearman’s rank correlation; prokaryotes: *r* = 0.035, *P* = 0.607; fungi: *r* = 0.006, *P* = 0.949; Fig. 5). Thus, the apparent scarcity of OTUs showing both habitat and host-plant preferences did not provide strong evidence for a trade-off between these two axes of specialization.

Plant taxa also differed in their association specificity for prokaryotic and fungal OTUs. For prokaryotes, significant plant-side association specificity was detected in five of the six host plant identities, except for *Enkianthus campanulatus* (Fig. 6). For fungi, significant plant-side association specificity was detected in all host plant identities (Fig. 7). Overall, *Rhododendron* spp., *Pinus parviflora*, and *Betula ermanii* showed consistently strong association specificity with both prokaryotic and fungal OTUs, whereas *Enkianthus campanulatus* showed a strong specificity signal only for fungal OTUs (Figs. 6–7). Representative specific plant–microbe associations included *Enkianthus campanulatus*–*Candidatus* Liberibacter [P_3583] (*z*-standardized 2DP score = 6.73; FDR < 0.01) and *Pinus parviflora*–*Silvibacterium* [P_0706] (*z*-standardized 2DP score = 5.87; FDR < 0.01) for prokaryotes (Fig. 6), and *Vaccinium smallii*–Cenangiaceae [F_0285] (*z*-standardized 2DP score = 5.10; FDR < 0.01) and *Betula ermanii*–*Hyaloscypha* [F_0074] (*z*-standardized 2DP score = 8.29; FDR < 0.01) for fungi (Fig. 7).

**Fig 6.**
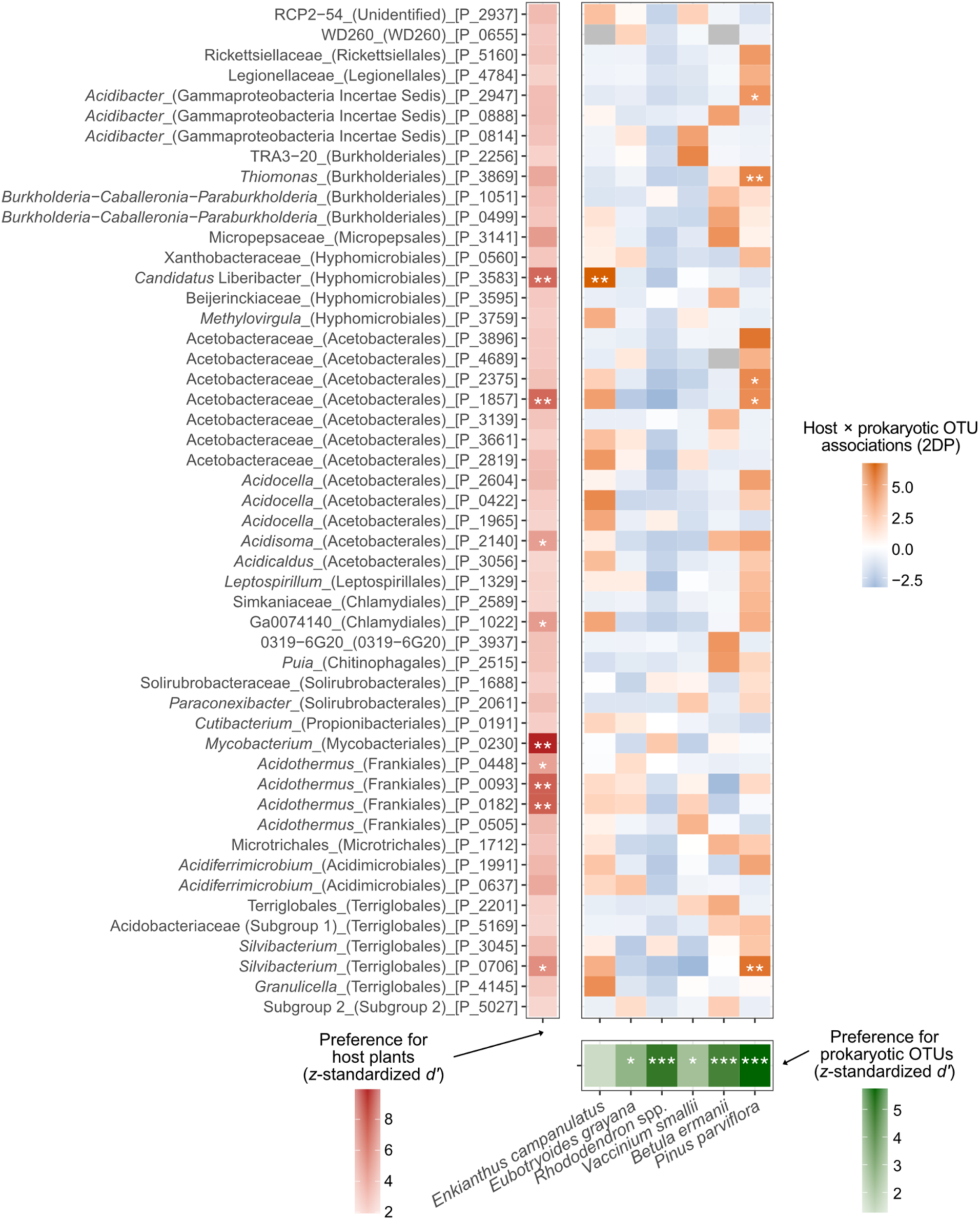
Preferences observed in plant–prokaryotic OTU associations. The top-50 prokaryotic OTUs with the largest absolute values of the *z*-standardized *d′* index of host plant preference, selected from the 222 OTUs detected from three or more root samples are shown. Likewise, the *z*-standardized *d′* index of plant-side association specificity for prokaryotic OTUs is shown for each host plant species. For each combination of prokaryotic OTU and plant species, the *z*-standardized two-dimensional preference index (2DP) indicates the specificity of the association. Statistical significance was assessed using two-tailed tests followed by false discovery rate (FDR) correction with the Benjamini–Hochberg method. Asterisks indicate significance levels: *, FDR < 0.05; **, FDR < 0.01; ***, FDR < 0.001. Grey cells indicate Z-scores that could not be calculated (NA); no significance symbols are shown for these cells.

**Fig 7.**
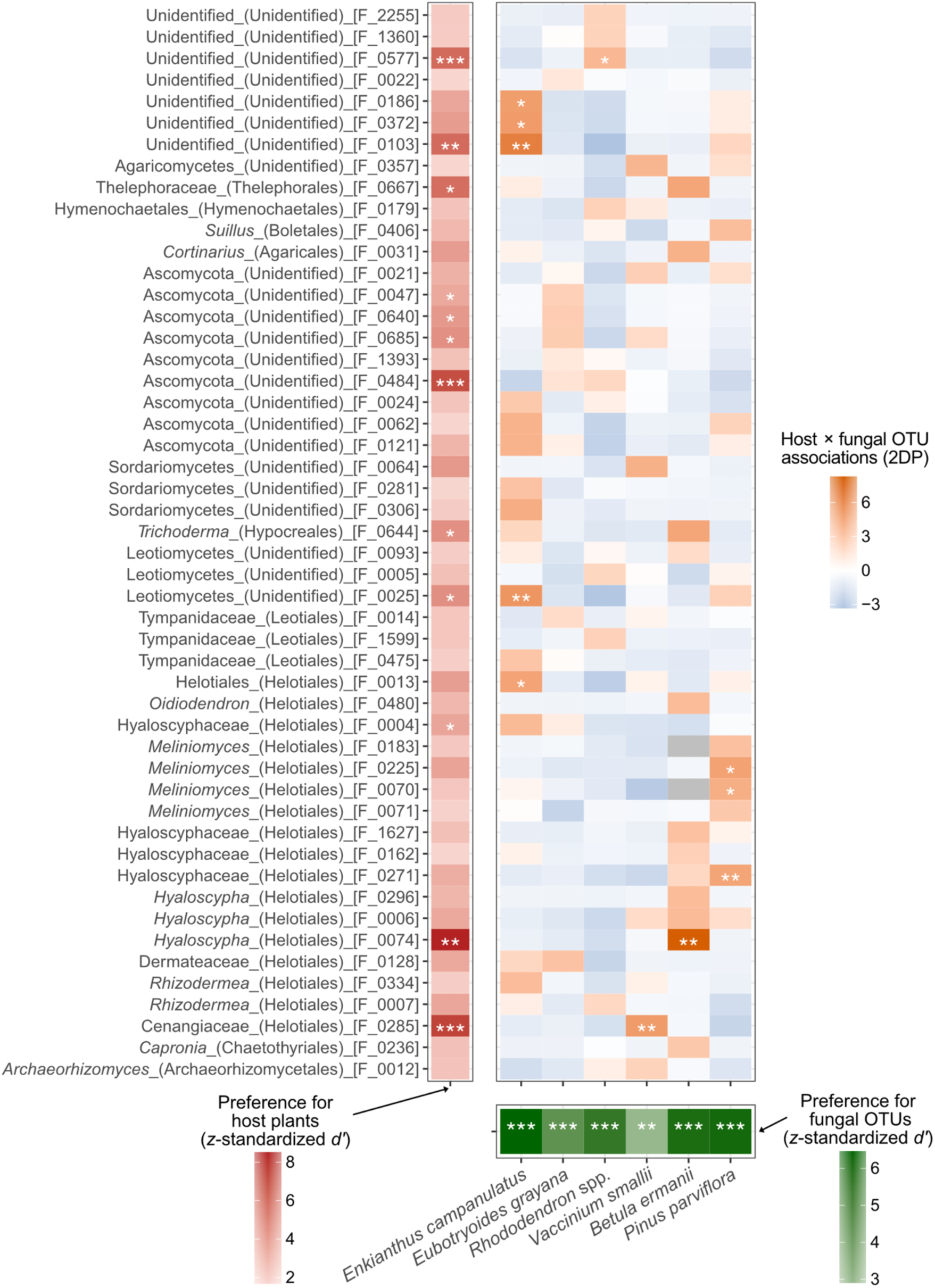
Preferences observed in plant–fungal OTU associations. The top-50 fungal OTUs with the largest absolute values of the *z*-standardized *d′* index of host plant preference, selected from the 137 OTUs detected from three or more root samples are shown. Likewise, the *z*-standardized *d′* index of plant-side association specificity for fungal OTUs is shown for each host plant species. For each combination of fungal OTU and plant species, the *z*-standardized two-dimensional preference index (2DP) indicates the specificity of the association. Statistical significance was assessed using two-tailed tests followed by false discovery rate (FDR) correction with the Benjamini–Hochberg method. Asterisks indicate significance levels: *, FDR < 0.05; **, FDR < 0.01; ***, FDR < 0.001. Grey cells indicate Z-scores that could not be calculated (NA); no significance symbols are shown for these cells.

### Phylogenetic signals in habitat and host preferences

To assess whether habitat and host preferences were phylogenetically conserved, we focused on the fungal family Hyaloscyphaceae, which exhibited substantial variation in both traits. Phylogenetic reconstruction using field-derived OTU sequences together with reference sequences resolved the sampled taxa into three major clades: Clade 1, including the reference database sequence of the ericoid mycorrhizal fungi *Rhizoscyphus ericae* [JQ711822]; Clade 2, including the reference database sequences of *Hyaloscypha variabilis* [OM238142] and *H. hepaticicola* [OM238143]; and Clade 3, including *Glutinomyces* database sequences [LC218288 and LC218304] (Fig. 8). The two preference traits varied substantially both within and among clades. Clade 1 generally exhibited weak habitat and host specialization, whereas Clade 2 included OTUs spanning a wide range of habitat and host preferences. Clade 3 contained several OTUs associated with forest-edge habitats. These patterns indicate substantial ecological heterogeneity within Hyaloscyphaceae but provide little evidence for phylogenetic conservatism of niche preferences.

**Fig 8.**
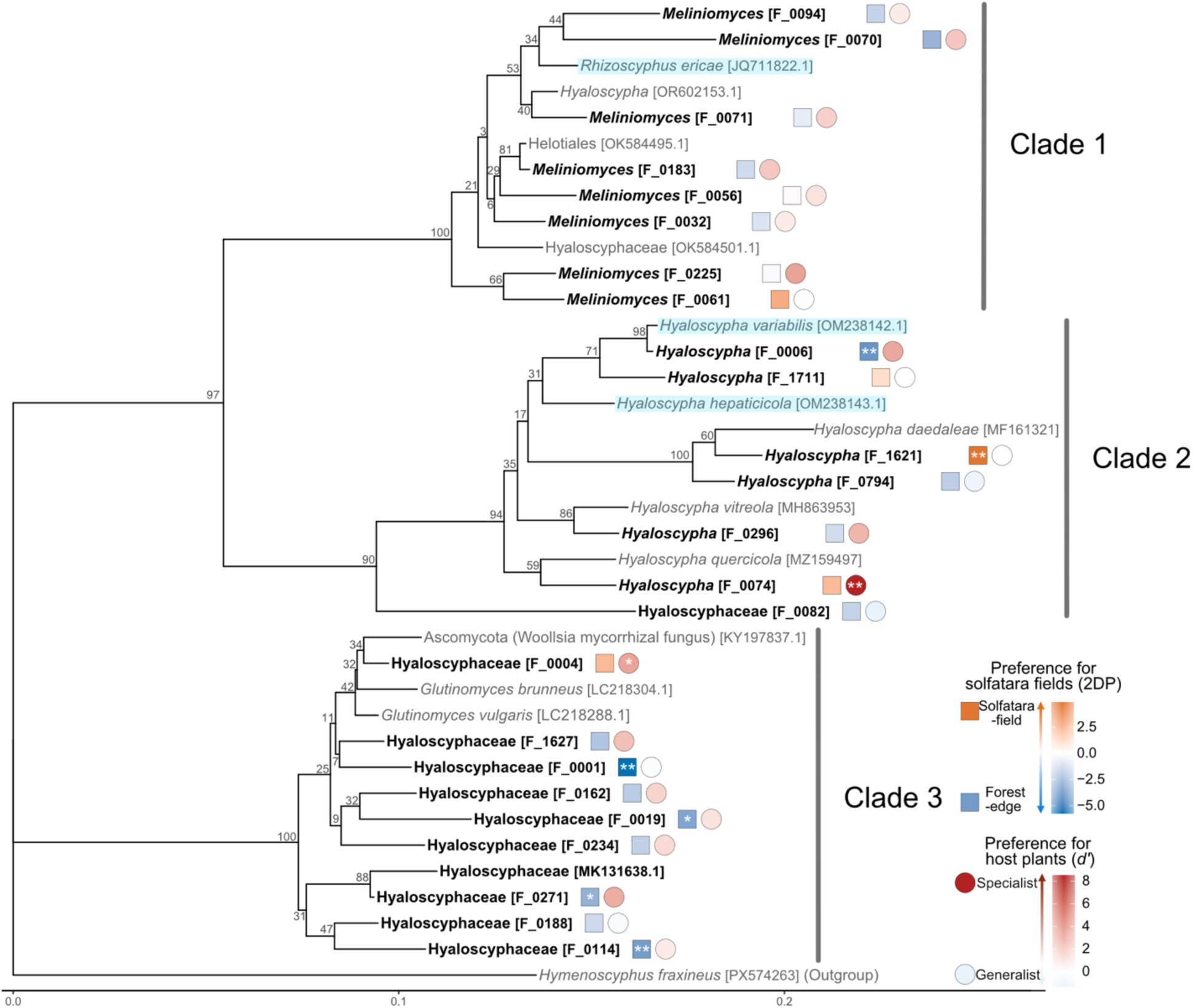
Distribution of habitat and host preferences across the Hyaloscyphaceae phylogenetic tree. The rooted phylogenetic tree was constructed using the neighbor-joining method based on the Kimura 2-parameter (K2P) model, with *Hymenoscyphus fraxineus* [PX574623] as the outgroup. Bootstrap values based on 1,000 replicates are indicated at the nodes. Tip labels are formatted as follows: bold text, OTUs obtained in this study; gray text, reference sequences; light blue highlighting, previously reported ericoid mycorrhizal fungi. Adjacent to each tip label, colored symbols represent two-dimensional preference index (2DP) estimates for solfatara-field preference (squares) and *z*-standardized *d′* estimates for host plant identity preference (circles). Asterisks indicate significance levels: *, FDR < 0.05; **, FDR < 0.01; ***, FDR < 0.001. *Rhizoscyphus* and *Meliniomyces* are currently classified as synonyms of *Hyaloscypha* (Fehrer et al., 2019).

Consistent with these observations, no significant phylogenetic signal was detected for habitat preference (*z*-standardized 2DP score; Blomberg’s *K* = 0.233, *P* = 0.609; Pagel’s *λ* = 0.071, *P* = 0.578). Likewise, host specialization (*z*-standardized *d′*) showed no detectable phylogenetic signal (Blomberg’s *K* = 0.288, *P* = 0.250; Pagel’s *λ* < 0.001, *P* = 1.000). These results provided no significant evidence for phylogenetic conservatism of habitat or host preferences across the sampled Hyaloscyphaceae lineages.

## Discussion

By focusing on contrasting habitats and dominant plant species in fumarole fields, we examined how environmental filtering and host plant identity collectively structure root-associated microbiomes under harsh environmental conditions. Our analyses identified prokaryotic and fungal OTUs that were strongly associated with either habitat (solfatara-field or forest-edge habitats) or particular host plants. However, only a few microbial OTUs showed significant associations with both habitat and host identity; instead, most specialized taxa were associated primarily with either a particular habitat or a particular host plant. Among the diverse microbial OTUs examined, several members of the fungal family Hyaloscyphaceae showed marked associations with specific habitats or host plants. These association patterns were not phylogenetically conserved within Hyaloscyphaceae, suggesting substantial niche differentiation within this ascomycete lineage. By combining OTU-level niche quantification with community-level analyses in a volcanic field system, this study provides insight into how plant-associated microbiomes assemble in extreme environments.

### Habitat and host effects on prokaryotic and fungal communities

By taking advantage of a natural system in which common host plant species occur across steep environmental gradients, we statistically partitioned the effects of abiotic environmental conditions and host plant identity on belowground microbiome structure. Both prokaryotic and fungal communities in plant root systems differed significantly among habitats and host identities (Fig. 4), indicating that root microbiome assembly is collectively shaped by environmental filtering and host-associated selection. Because root-associated microbes are recruited from the microbial species pool present in the surrounding soil [12, 80, 81], differences in soil communities between habitats are expected to influence the initial pool available for root colonization. Consistent with this conceptual framework, bulk soil communities differed markedly between solfatara-field and forest-edge habitats, indicating environmental filtering of the habitat-specific microbial pool (Fig. S5; Table S8). These results support a sequential assembly process in which environmental filtering first shapes the available soil microbial pool, after which host-associated selection further filters this pool to establish the root microbiomes.

Patterns of *β*−diversity further reinforced this concept. Root-associated prokaryotic and fungal communities differed significantly according to both habitat and host identity (Table S4). Moreover, root communities exhibited greater within-group dispersion than bulk soil communities (Fig. S6; Table S10), suggesting that host-specific selection generates greater compositional heterogeneity than environmental filtering alone. Because soils represent the common microbial source pool, whereas individual host species selectively recruit subsets of microbes, increased dispersion among root communities likely reflects differences in host-specific symbiont recruitment rather than stochastic variation.

Although habitat and host identity significantly structured root-associated microbial communities, they explained only a limited proportion of the total community variation (Table S4). This is not unexpected in field-based microbiome studies, in which community composition is influenced by many factors that are difficult to measure explicitly, including fine-scale variation in soil chemistry, microclimate, spatial structure, plant physiological status, microbe–microbe interactions, historical contingency, and stochastic processes such as ecological drift [11, 82, 83]. Thus, the unexplained variation in our statistical analyses does not negate the roles of environmental filtering and host-associated selection but rather indicates that these processes operate together with additional ecological and historical factors. Taken together with the consistent differentiation observed in community composition and diversity, our results suggest that environmental filtering and host-associated selection are two important processes shaping root microbiome assembly in an extreme environment.

### Niche differentiation patterns at the microbial OTU level

To understand how niche differentiation contributes to plant microbiome assembly, we quantified habitat and host-plant associations for individual prokaryotic and fungal OTUs. A prevailing conceptual model proposes that plant-associated microbiomes are assembled through hierarchical filtering, whereby environmental conditions first determine the pool of available microbes and host plants subsequently recruit subsets of this pool through physiological and immunological selection [12, 80, 81]. Based on this framework, we hypothesized that the environmental contrast between solfatara-field and forest-edge habitats, together with host selection by Ericaceae and ectomycorrhizal plants, would result in the prevalence of microbes specialized for both environmental and host niches.

Unexpectedly, OTUs showing significant specialization along both habitat and host-plant axes were rare (Fig. 5). Instead, most specialized OTUs were associated with either a particular habitat or a particular host plant. This pattern suggests that community-level differentiation can emerge from the combined contributions of taxa specialized along different niche axes, rather than from a large number of dual-specialist taxa. Despite this trend, no significant trade-off was detected between habitat and host-plant associations. Therefore, the rarity of dual specialists should not be interpreted as evidence for a simple negative relationship between these two axes of specialization. Rather, our results indicate that habitat specialization and host-plant associations represent largely independent dimensions of microbial niche differentiation in fumarole fields.

Among root-associated microbes specialized along a single niche axis, fungal OTUs showing significant associations with contrasting habitats are particularly relevant to the concept of habitat-adapted symbiosis, in which microbial symbionts derived from specific environments can enhance plant tolerance to local stress conditions [6]. In our analysis, the presence of solfatara-field-specific OTUs was observed in fungal but not in prokaryotic communities (Fig. 5), highlighting the potential roles of specific fungi for mediating plant establishment under fumarolic environmental gradients.

Among the fungal taxa highlighted by our analysis, members of the family Hyaloscyphaceae were of particular interest because closely related OTUs were detected across both contrasting habitats (Fig. 5). Members of Hyaloscyphaceae are known to exhibit high ecological diversity, ranging from saprotrophic and root-endophytic lifestyles to mutualistic ericoid and ectomycorrhizal associations, some of which have been reported to enhance plant nutrient acquisition and stress tolerance [84–89]. Moreover, the presence of Hyaloscyphaceae fungi can modify the community assembly of other root-associated fungi, thereby potentially organizing the entire microbiome functions in plant root systems [90]. The occurrence patterns of Hyaloscyphaceae OTUs in this study may reflect their high ecological versatility and pivotal functional roles in plant-root interactions. Among the Hyaloscyphaceae fungi detected in our analysis, *Hyaloscypha* [F_1621], a solfatara-field-associated OTU, was closely related to *Hyaloscypha daedaleae*, a saprobic species whose close relatives can form ericoid mycorrhizal associations and promote the growth of Ericaceae plants [91]. This fact raises the possibility that the *Hyaloscypha* lineage supports host nutrition by decomposing scarce organic substrates in nutrient-poor solfatara soils. In contrast, Hyaloscyphaceae [F_0001], which was associated with roots in forest-edge habitats, is phylogenetically closest to *Glutinomyces vulgaris*, a root-endophytic lineage frequently isolated from temperate forest ecosystems [92], suggesting potential ecological diversification within this fungal family. Likewise, *Hyaloscypha* [F_0006], which is closely related to the ericoid mycorrhizal fungus *Hyaloscypha variabilis* [85, 86], may contribute to host tolerance in acidic, nutrient-poor environments. Although these functional interpretations remain speculative, the contrasting environmental associations of closely related Hyaloscyphaceae OTUs identify this lineage as a promising target for future studies testing the ecological roles of these fungi in plant establishment and persistence in fumarolic environments.

### Phylogenetic conservatism of niche preference in Hyaloscyphaceae

We further examined whether habitat and host preference traits were phylogenetically conserved within Hyaloscyphaceae. Although phylogenetic niche conservatism has been widely reported for various traits in both prokaryotes and fungi [93–95], neither habitat specialization nor host-plant association exhibited a significant phylogenetic signal within this fungal family. Closely related Hyaloscyphaceae OTUs frequently differed in their associations with solfatara-field habitats, forest-edge habitats, or host plants (Fig. 8), suggesting that habitat and host preference traits may be evolutionarily labile among closely related taxa. Such lability may facilitate ecological differentiation across contrasting habitats and host plants. However, this interpretation should be treated cautiously because our phylogenetic analysis was restricted to a single fungal family selected for its pronounced variation in habitat and host associations. Therefore, whether weak phylogenetic constraint is a broader feature of root-associated fungi or a lineage-specific property of Hyaloscyphaceae remains to be tested.

### Conclusions

This study indicates that root microbiome assembly in dominant Ericaceae and ectomycorrhizal plants inhabiting fumarole fields is collectively shaped by environmental filtering and host identity. Notably, differentiation of root-associated microbial communities across these niche axes arose primarily through the collective assembly of microbes specialized along a single niche axis, rather than taxa simultaneously specialized to both environmental conditions and host plants. These findings suggest that complex community-level patterns can emerge from the combined contributions of microbes with distinct ecological preferences. With respect to potential contributions to plant adaptation to harsh volcanic environments, Hyaloscyphaceae fungi represent particularly promising candidates for understanding microbiome-mediated plant adaptation to fumarolic environments. Their contrasting habitat associations despite low host specificity, together with the weak phylogenetic signal detected for habitat and host preference traits, suggest considerable ecological and evolutionary flexibility within this fungal lineage.

Our findings provide a snapshot of a dynamic ecological process. Root-associated microbiomes are likely to vary across spatial scales, seasons, and host developmental stages, and a more comprehensive understanding of microbiome assembly will require long-term and geographically replicated studies. In addition, DNA metabarcoding reveals community composition and ecological associations but cannot directly determine microbial activity or functions. Future studies integrating metagenomics, transcriptomics, microbial isolation, and inoculation experiments will be essential for establishing causal links among microbial occurrence, microbial function, and host physiological responses, ultimately advancing our understanding of how root-associated microbes contribute to plant persistence and adaptation in extreme volcanic environments.

## Abbreviations

a.s.l.: above sea level
FDR: False discovery rate
ITS: Internal transcribed spacer
NCBI: National Center for Biotechnology Information
OTU: Operational taxonomic unit
PCA: Principal component analysis
PCR: Polymerase chain reaction
PERMANOVA: Permutational analysis of variance
PERMDISP: Permutational analysis of dispersion
PCoA: Principal coordinate analysis
rRNA: ribosomal RNA

## Acknowledgements

We thank the Miyagi Prefectural Northern Forest Management Office and the Miyagi Prefectural Northern Regional Development Office for granting permission to conduct field surveys at Mt. Arao. We are also grateful to Yoshihisa Suyama (Tohoku University) for his generous support and for providing access to research facilities that made this fieldwork possible.

## Author contributions

AM, MN, and HT designed the work. AM, MN, IH, and HT performed fieldwork. AM performed the molecular experiments. AM analyzed the data. AM and HT wrote the manuscript based on discussion with all the co-authors.

## Funding

This work was conducted based on the financial support by JST FOREST (JPMJFR2048), JST CREST (JPMJCR23N5), and Kyoto University CeLiSIS Program (23CeLiSIS-02) to HT and Kyoto University SPRING Program (A94261500036) to AM.

## Data availability

The sequencing data of the prokaryotic 16S rRNA gene, the fungal ITS, and the plant ITS regions are available from the DNA Data Bank of Japan (DDBJ) with the accession number PRJDB42913 [to be released after the acceptance of this manuscript].

## Code availability statement

All computational codes used to analyze the data are available at the zenodo repository at https://doi.org/10.5281/zenodo.22955378.

## Declarations

### Ethics approval and consent to participate

Not applicable.

### Consent for publication

Not applicable.

### Generative AI statement

ChatGPT (OpenAI) and Gemini (Google) were used to assist with coding and language editing. The authors conducted all analyses and scientific interpretations and reviewed and verified the final manuscript text.

### Competing interests

H.T. is a founder, director, and shareholder of Sunlit Seedlings Ltd., a Kyoto University spin-off, which had no role in this study. The other authors declare no competing interests.

## Notes

https://doi.org/10.5281/zenodo.22955378

