## Supplementary Figures and Tables for "Environmental filtering and host identity collectively shape root-associated microbiomes of Ericaceae and ectomycorrhizal plants in fumarole fields"

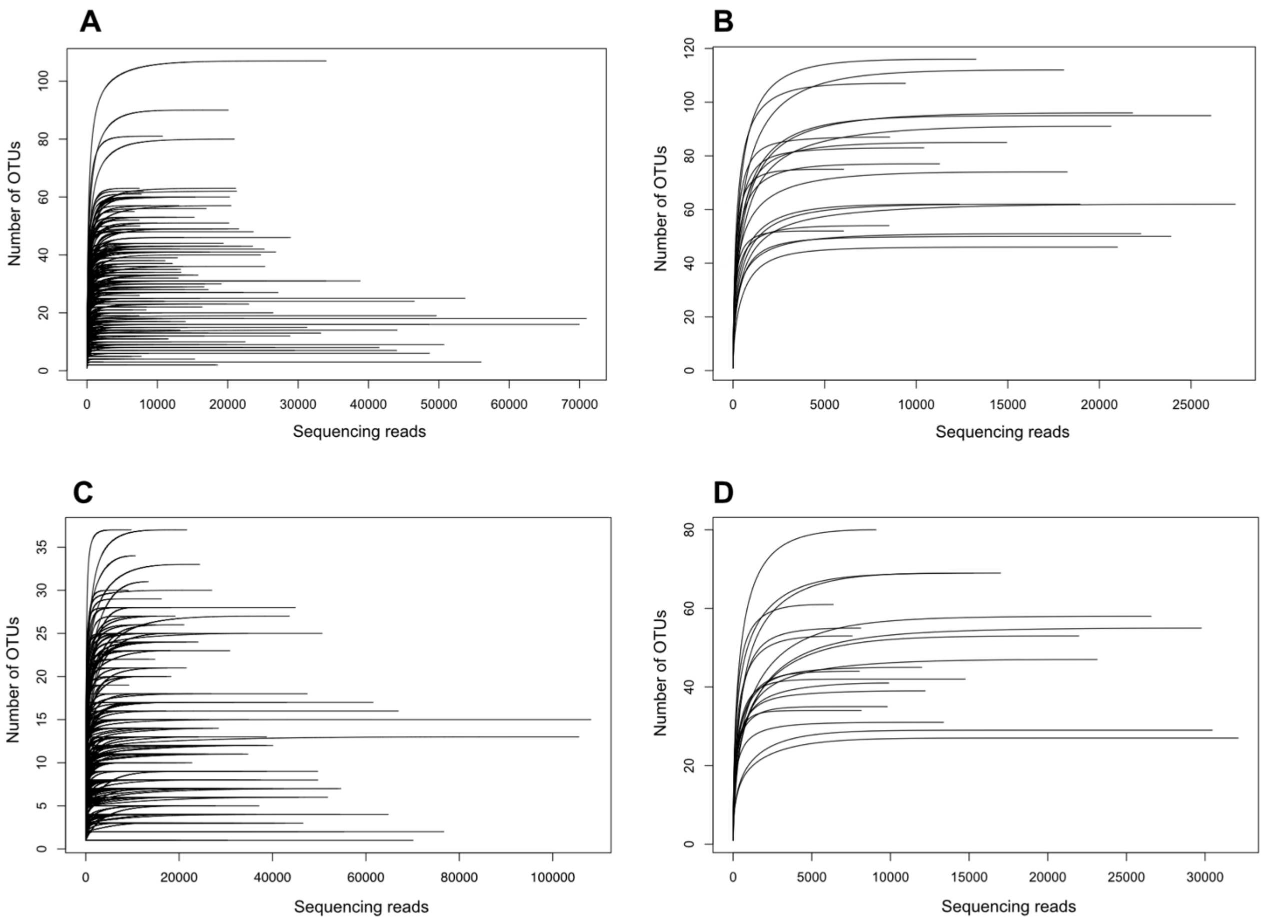


**Fig S1. Rarefaction curves for prokaryotic and fungal communities.**
Rarefaction curves are shown for prokaryotic communities in (A) root and (B) soil samples and for fungal communities in (C) root and (D) soil samples. The horizontal and vertical axes indicate the number of sequenced reads and the observed number of operational taxonomic units (OTUs), respectively. Samples were included in downstream analyses only when they had at least 2,000 reads for root-associated samples or 5,000 reads for soil samples. For downstream analyses, each sample was rarefied to the minimum sequencing depth.

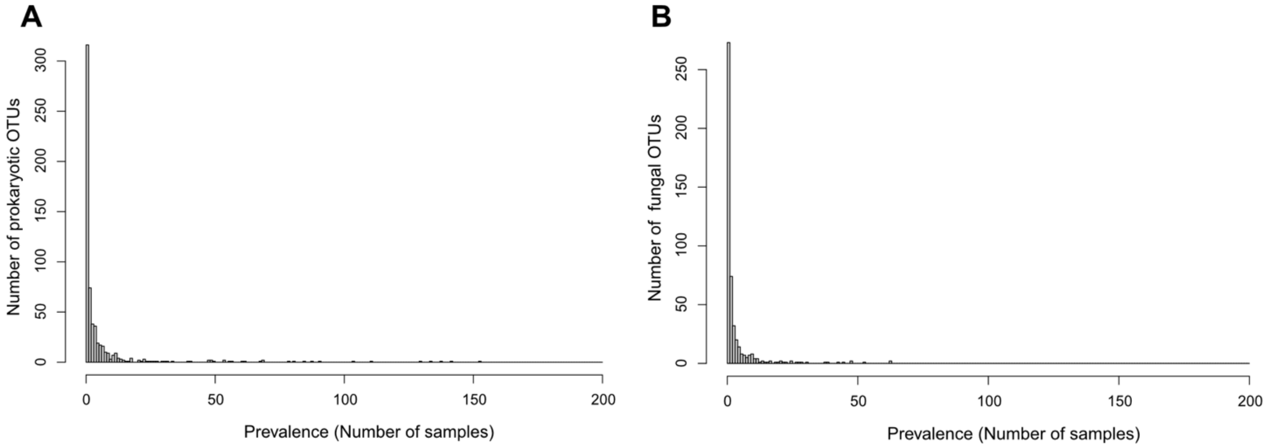


**Fig S2. Prevalence distributions of OTUs in root-associated prokaryotic and fungal communities.**

Histograms show the prevalence (occurrence) distributions of OTUs in root-associated (A) prokaryotic and (B) fungal communities. The horizontal axis indicates the number of samples in which each OTU was detected, and the vertical axis indicates the number of OTUs. Histograms were plotted with a bin width of one sample.

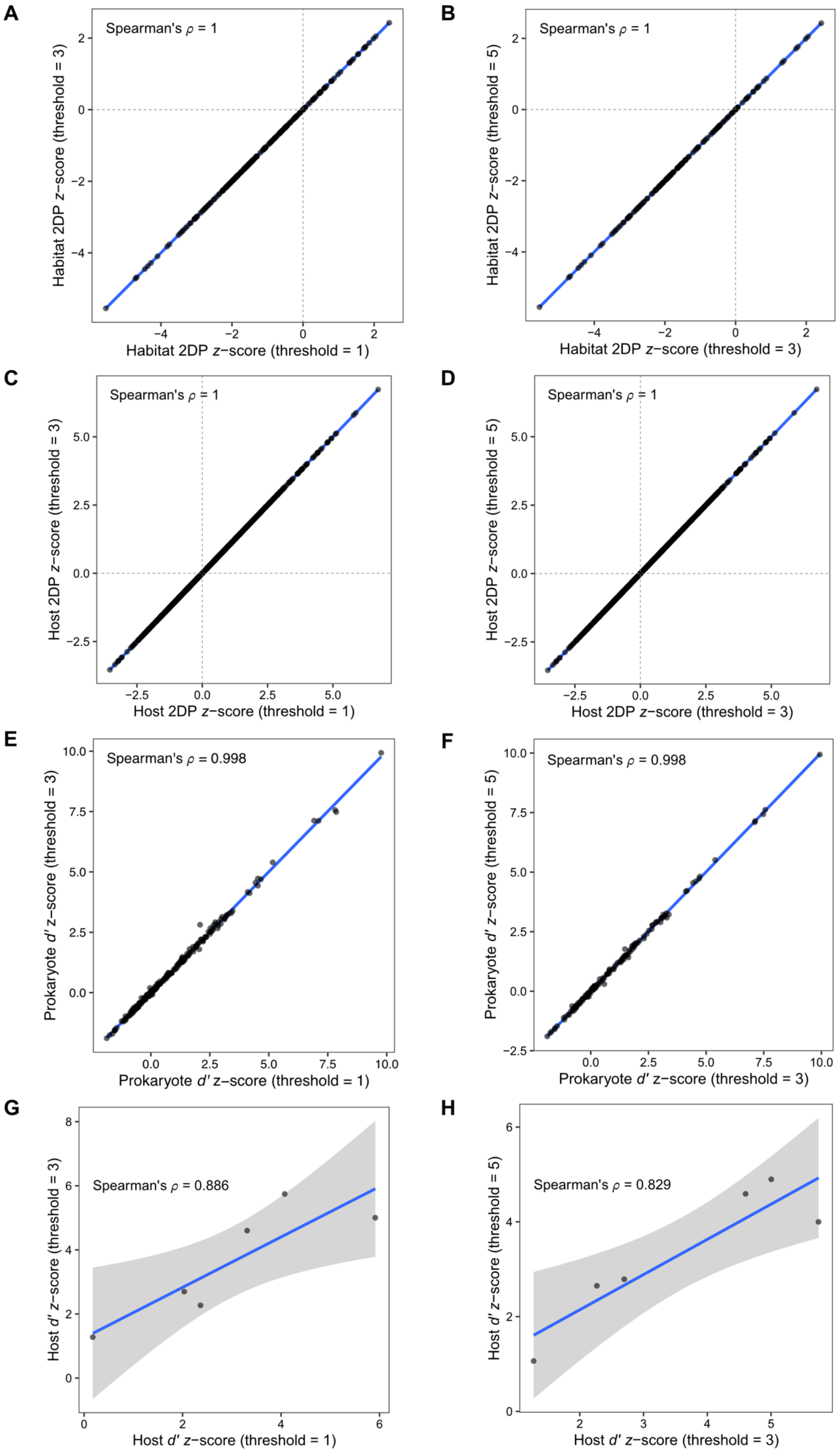

A
Spearman's ρ = 1
Habitat 2DP z−score (threshold = 3)
Habitat 2DP z−score (threshold = 1)
B
Spearman's ρ = 1
Habitat 2DP z−score (threshold = 5)
Habitat 2DP z−score (threshold = 3)
C
Spearman's ρ = 1
Host 2DP z−score (threshold = 3)
Host 2DP z−score (threshold = 1)
D
Spearman's ρ = 1
Host 2DP z−score (threshold = 5)
Host 2DP z−score (threshold = 3)
E
Spearman's ρ = 0.998
Prokaryote d′ z−score (threshold = 3)
Prokaryote d′ z−score (threshold = 1)
F
Spearman's ρ = 0.998
Prokaryote d′ z−score (threshold = 5)
Prokaryote d′ z−score (threshold = 3)
G
Spearman's ρ = 0.886
Host d′ z−score (threshold = 3)
Host d′ z−score (threshold = 1)
H
Spearman's ρ = 0.829
Host d′ z−score (threshold = 5)
Host d′ z−score (threshold = 3)

**Fig S3. Consistency of *z*-scores across prevalence thresholds in 2DP and *d′* randomization analyses of prokaryotic communities.**

Robustness of *z*-scores across prevalence (occurrence) thresholds was evaluated by pairwise comparisons between thresholds of 1 and 3 samples and between thresholds of 3 and 5 samples per prokaryotic OTU. The blue line represents a linear regression line; points tighter around this line indicate stronger monotonic agreement between thresholds. Spearman's rank correlation coefficient $\rho$ is shown in the upper left corner of each panel. Comparisons are shown for prokaryotic OTU–habitat 2DP scores [(A) threshold 1 vs. threshold 3; (B) threshold 3 vs. threshold 5], prokaryotic OTU–host plant 2DP scores [(C) threshold 1 vs. threshold 3; (D) threshold 3 vs. threshold 5], prokaryotic OTU-side *d′* host preference scores [(E) threshold 1 vs. threshold 3; (F) threshold 3 vs. threshold 5], and host-side *d′* association specificity for prokaryotic OTUs [(G) threshold 1 vs. threshold 3; (H) threshold 3 vs. threshold 5].

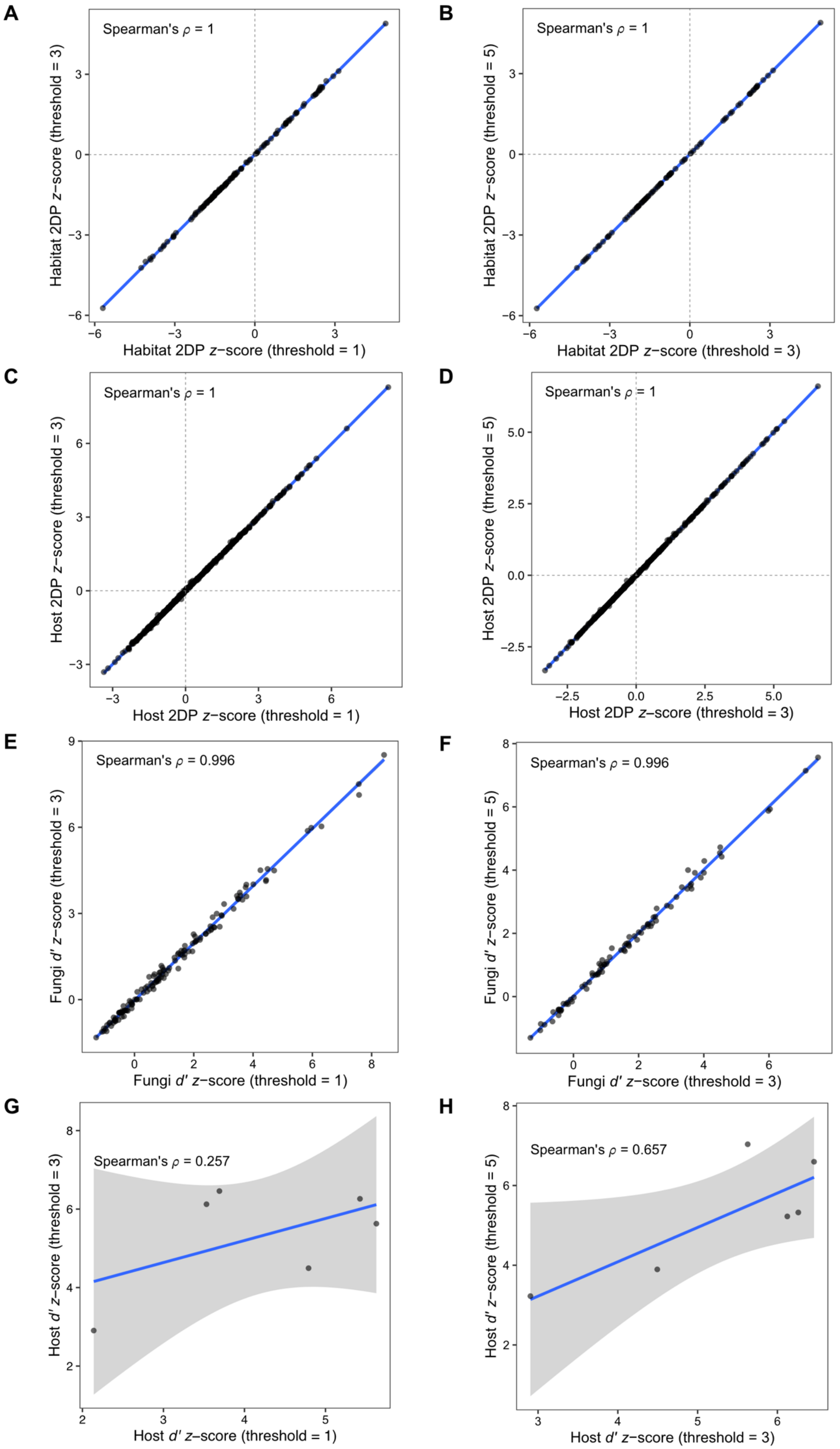
A
Spearman's ρ = 1
Habitat 2DP z−score (threshold = 3)
Habitat 2DP z−score (threshold = 1)
B
Spearman's ρ = 1
Habitat 2DP z−score (threshold = 5)
Habitat 2DP z−score (threshold = 3)
C
Spearman's ρ = 1
Host 2DP z−score (threshold = 3)
Host 2DP z−score (threshold = 1)
D
Spearman's ρ = 1
Host 2DP z−score (threshold = 5)
Host 2DP z−score (threshold = 3)
E
Spearman's ρ = 0.996
Fungi d′ z−score (threshold = 3)
Fungi d′ z−score (threshold = 1)
F
Spearman's ρ = 0.996
Fungi d′ z−score (threshold = 5)
Fungi d′ z−score (threshold = 3)
G
Spearman's ρ = 0.257
Host d′ z−score (threshold = 3)
Host d′ z−score (threshold = 1)
H
Spearman's ρ = 0.657
Host d′ z−score (threshold = 5)
Host d′ z−score (threshold = 3)

**Fig S4. Consistency of *z*-scores across prevalence thresholds in 2DP and *d′* randomization analyses of fungal communities.**

Robustness of *z*-scores across prevalence (occurrence) thresholds was evaluated by pairwise comparisons between thresholds of 1 and 3 samples and between thresholds of 3 and 5 samples per fungal OTU. The blue line represents a linear regression line; points tighter around this line indicate stronger monotonic agreement between thresholds. Spearman's rank correlation coefficient $\rho$ is shown in the upper left corner of each panel. Comparisons are shown for fungal OTU–habitat 2DP scores [(A) threshold 1 vs. threshold 3; (B) threshold 3 vs. threshold 5], fungal OTU–host plant 2DP scores [(C) threshold 1 vs. threshold 3; (D) threshold 3 vs. threshold 5], fungal OTU-side *d′* host preference scores [(E) threshold 1 vs. threshold 3; (F) threshold 3 vs. threshold 5], and host-side *d′* association specificity for fungal OTUs [(G) threshold 1 vs. threshold 3; (H) threshold 3 vs. threshold 5].

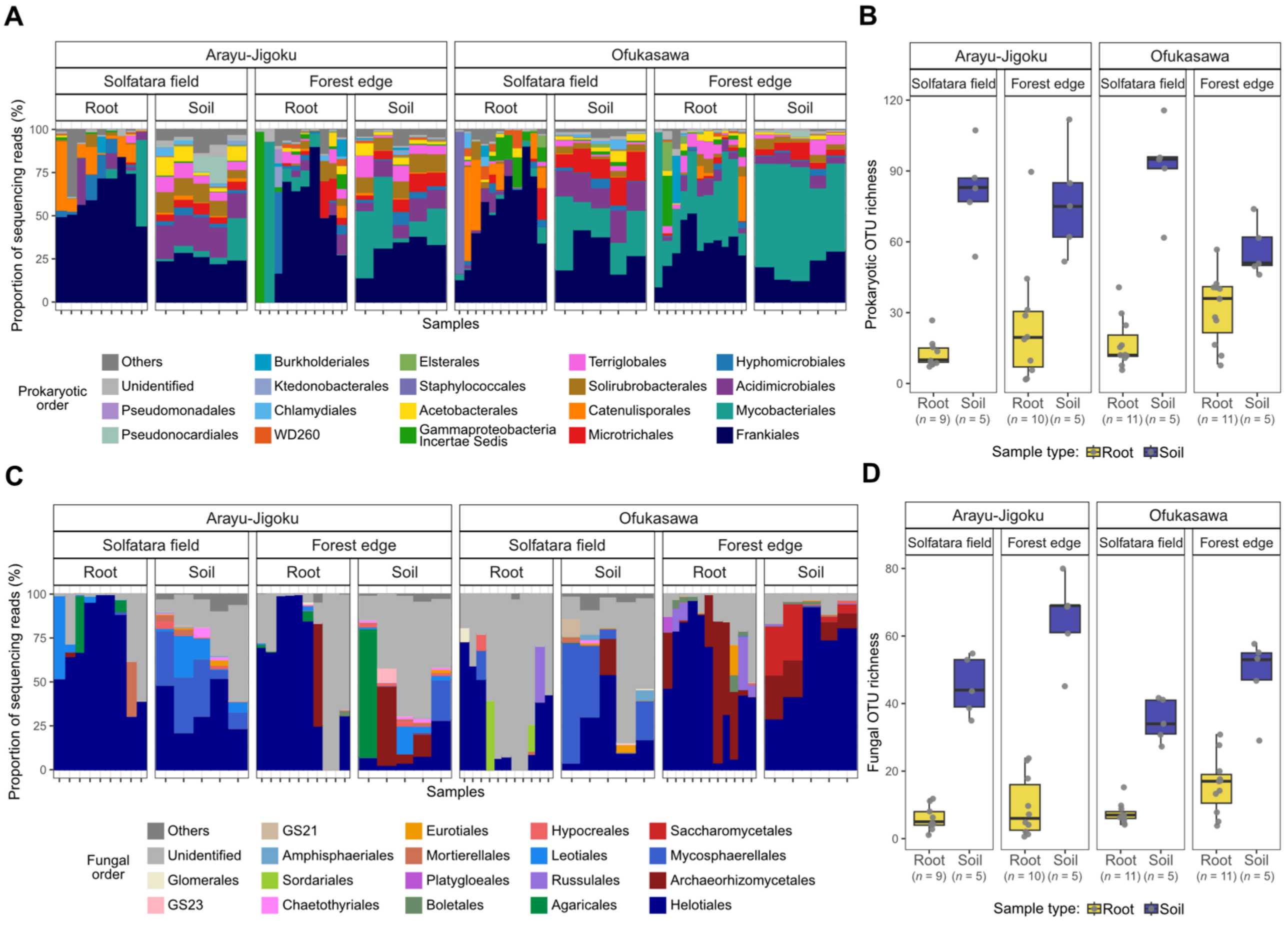


**Fig S5. Prokaryotic and fungal community composition and OTU richness in soil and root samples from representative sampling points.**

(A, B) Order-level taxonomic composition of prokaryotic (A) and fungal (B) communities in soil and root samples. Samples were ordered by hierarchical clustering using the average linkage method based on Bray–Curtis dissimilarity.

(C, D) Comparison of prokaryotic (C) and fungal (D) OTU richness between soil and root samples from solfatara-field and forest-edge habitats at each study site. Soil OTU richness was significantly higher than root OTU richness for both prokaryotes and fungi (GLMM, $P < 0.001$, for both).

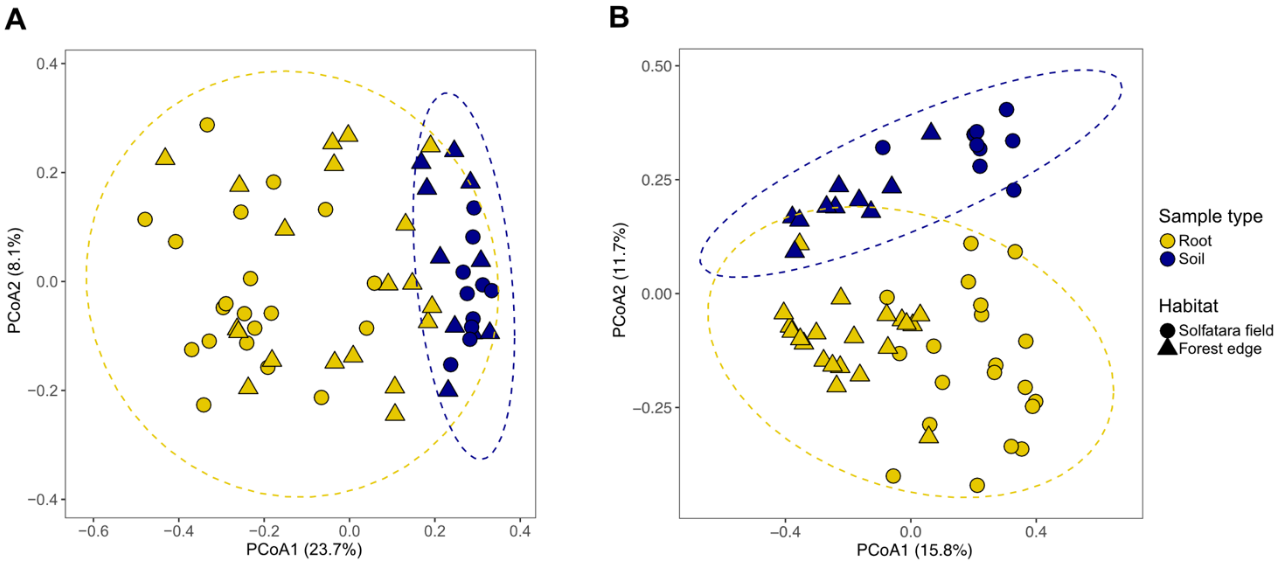


**Fig S6. Variation in root-associated prokaryotic and fungal community structure across sample types and habitats.**

PCoA plots showing variation in prokaryotic (A) and fungal (B) community composition based on the Sørensen dissimilarity index for representative root and soil samples from solfatara-field and forest-edge habitats. Points represent individual samples, colored by sample type (root or soil) and shaped by habitats (solfatara-field or forest-edge habitats). Ellipses indicate 95% confidence intervals for each habitat (red, solfatara field; blue, forest edge).

**Table S1. Factor loadings of soil chemical properties for the first nine principal components.** Factor loadings are shown for principal components 1–9 (PC1–PC9).

| Factors | PC1 | PC2 | PC3 | PC4 | PC5 | PC6 | PC7 | PC8 | PC9 |
|---|---|---|---|---|---|---|---|---|---|
| $Na^{+}$ | 0.380 | -0.089 | -0.381 | 0.322 | 0.132 | -0.257 | -0.283 | -0.535 | -0.387 |
| $K^{+}$ | 0.436 | -0.026 | -0.216 | 0.118 | 0.193 | 0.053 | 0.057 | -0.074 | 0.836 |
| $Ca^{2+}$ | 0.333 | 0.324 | 0.287 | 0.219 | -0.598 | 0.149 | 0.427 | -0.300 | -0.048 |
| $Mg^{2+}$ | 0.419 | 0.007 | -0.218 | 0.156 | 0.266 | 0.044 | 0.466 | 0.589 | -0.340 |
| $Zn^{2+}$ | 0.363 | -0.001 | 0.454 | -0.017 | -0.155 | -0.645 | -0.332 | 0.329 | 0.059 |
| $Pb^{2+}$ | 0.269 | -0.357 | 0.307 | -0.621 | 0.284 | -0.040 | 0.326 | -0.350 | -0.101 |
| $Mn^{2+}$ | 0.078 | 0.725 | 0.309 | -0.020 | 0.544 | 0.168 | -0.181 | -0.099 | -0.068 |
| $Al^{3+}$ | 0.393 | -0.199 | 0.065 | -0.158 | -0.194 | 0.658 | -0.515 | 0.166 | -0.119 |
| pH | -0.118 | -0.440 | 0.528 | 0.633 | 0.281 | 0.171 | 0.047 | -0.052 | 0.012 |

**Table S2. Results of permutational multivariate analysis of variance (PERMANOVA) testing for differences in soil chemical composition between habitats.**

The table reports degrees of freedom (*df*), sums of squares (SS), proportion of variance explained ($R^2$), $F$ statistics ($F$), and permutation-based $P$ values ($P$) for each term. $R^2$ represents the fraction of total variation in soil chemical composition explained by habitat grouping.

| Factors | *df* | SS | $R^2$ | $F$ | $P$ |
|---|---|---|---|---|---|
| Habitat | 1 | 60.513 | 0.280 | 8.951 | < 0.001 |
| Residual | 23 | 155.487 | 0.720 | NA | |
| Total | 24 | 216.000 | 1.000 | NA | |

**Table S3. Results of permutational analysis of multivariate dispersions (PERMDISP) testing for differences in within-group dispersion of soil chemical composition between habitats.** The table reports degrees of freedom (*df*), sums of squares (SS), mean squares (MS), *F* statistics (*F*), and permutation-based *P* values (*P*) for each term. Significant results indicate heterogeneity in multivariate dispersion, that is, differences in within-habitat variability of soil chemical composition.

| **Factors** | ***df*** | **SS** | **MS** | ***F*** | ***P*** |
|---|---|---|---|---|---|
| Habitat | 1 | 19.353 | 19.353 | 24.416 | < 0.001 |
| Residuals | 23 | 18.230 | 0.793 | | |

**Table S4. Results of PERMANOVA testing for differences in root-associated community composition of (A) prokaryotes and (B) fungi.**

PERMANOVA was performed based on Sørensen dissimilarity, with habitat, host plant identity, and their interaction included as explanatory variables. "Overall model" indicates the PERMANOVA including all three terms, and "Within site" indicates that permutations were restricted within study sites. The table reports degrees of freedom (*df*), sums of squares (SS), proportion of variance explained ($R^2$), F statistics (*F*), and permutation-based *P* values (*P*) for each term. $R^2$ represents the proportion of variation explained by each term, calculated sequentially according to the model structure. "Habitat × Host" denotes the interaction between habitat and host plant identity.

(A)

| **Test** | **Permutation restriction** | **Factor** | ***df*** | **SS** | $R^2$ | ***F*** | ***P*** |
|---|---|---|---|---|---|---|---|
| Overall model | Within site | Habitat | 1 | 3.043 | 0.052 | 11.883 | < 0.001 |
| | | Host plant | 5 | 3.845 | 0.066 | 3.003 | < 0.001 |
| | | Habitat × Host | 5 | 1.639 | 0.028 | 1.280 | 0.085 |
| | | Residual | 194 | 49.674 | 0.853 | | |
| | | Total | 205 | 58.201 | 1.000 | | |

(B)

| **Test** | **Permutation restriction** | **Factor** | ***df*** | **SS** | $R^2$ | ***F*** | ***P*** |
|---|---|---|---|---|---|---|---|
| Overall model | Within site | Habitat | 1 | 5.547 | 0.066 | 15.444 | < 0.001 |
| | | Host plant | 5 | 6.929 | 0.083 | 3.859 | < 0.001 |
| | | Habitat × Host | 5 | 2.461 | 0.029 | 1.371 | 0.021 |
| | | Residual | 192 | 68.955 | 0.822 | | |
| | | Total | 203 | 83.893 | 1.000 | | |

**Table S5. Results of variation partitioning analysis for root-associated prokaryotic and fungal community composition.**

Variation partitioning was performed based on Sørensen dissimilarity for (A) prokaryotic and (B) fungal communities to quantify the fractions of variation attributable to habitat and host plant identity. Fractions are expressed as adjusted $R^2$ values, accounting for model complexity. "Habitat + Host" represents the variation explained by the full model containing both variables. "Habitat (unique)" and "Host (unique)" represent the fractions uniquely attributable to each variable after accounting for the other, whereas "Shared (Habitat ∩ Host)" represents their joint fraction. "Habitat (total)" and "Host (total)" represent the variation explained by each variable without accounting for the other.

(A)

| **Partition** | **Adjusted $R^2$** | ***df*** |
|---|---|---|
| Habitat (total) | 0.048 | 1 |
| Host (total) | 0.055 | 5 |
| Habitat+Host | 0.092 | 6 |
| Habitat (unique) | 0.037 | 1 |
| Host (unique) | 0.044 | 5 |
| Shared (Habitat ∩ Host) | 0.011 | 0 |
| Residual | 0.908 | |

(B)

| **Partition** | **Adjusted $R^2$** | ***df*** |
|---|---|---|
| Habitat (total) | 0.061 | 1 |
| Host (total) | 0.075 | 5 |
| Habitat+Host | 0.123 | 6 |
| Habitat (unique) | 0.048 | 1 |
| Host (unique) | 0.061 | 5 |
| Shared (Habitat ∩ Host) | 0.013 | 0 |
| Residual | 0.877 | |

**Table S6. Results of PERMDISP testing for differences in within-group dispersion of root-associated community composition among habitats and host plant identities.**

PERMDISP was performed separately for habitat and host plant identity for (A) prokaryotic and (B) fungal communities based on Sørensen dissimilarity, with permutations restricted within study sites for both tests. "Habitat" and "Host plant" in the Test column indicate the grouping variable used in each separate PERMDISP analysis. Significant results indicate heterogeneity in multivariate dispersion, that is, differences in within-habitat or within-host variation in community composition. The table reports degrees of freedom (*df*), sums of squares (SS), mean squares (MS), *F* statistics (*F*), and permutation-based *P* values (*P*).

(A)

| Test | Permutation restriction | Factor | *df* | SS | MS | *F* | *P* |
|---|---|---|---|---|---|---|---|
| Habitat | Within site | Habitat | 1 | 0.030 | 0.030 | 2.249 | 0.134 |
| | | Residual | 204 | 2.704 | 0.013 | | |
| Host plant | | Host plant | 5 | 0.321 | 0.064 | 5.045 | < 0.001 |
| | | Residual | 200 | 2.546 | 0.013 | | |

(B)

| Test | Permutation restriction | Factor | *df* | SS | MS | *F* | *P* |
|---|---|---|---|---|---|---|---|
| Habitat | Within site | Habitat | 1 | 0.001 | 0.001 | 0.212 | 0.632 |
| | | Residual | 202 | 1.162 | 0.006 | | |
| Host plant | | Host plant | 5 | 0.200 | 0.040 | 4.737 | 0.001 |
| | | Residual | 198 | 1.669 | 0.008 | | |

**Table S7. Pairwise PERMDISP comparisons of root-associated community dispersion among host plant identities.**

Pairwise comparisons were performed for (A) prokaryotic and (B) fungal communities based on Sørensen dissimilarity, with permutations restricted within study sites. The table reports *P* values adjusted for multiple comparisons using the Benjamini–Hochberg false discovery rate (FDR) method. Significant FDR-adjusted *P* values indicate differences in multivariate dispersion, that is, differences in within-host variation in community composition, between host plant identities.

(A)

| Host comparison | Permutation restriction | FDR |
|---|---|---|
| *Betula ermanii–Enkianthus campanulatus* | Within site | 0.559 |
| *Betula ermanii–Eubotryoides grayana* | Within site | 0.892 |
| *Betula ermanii–Pinus parviflora* | Within site | 0.455 |
| *Betula ermanii–Rhododendron spp.* | Within site | 0.559 |
| *Betula ermanii–Vaccinium smallii* | Within site | 0.455 |
| *Enkianthus campanulatus–Eubotryoides grayana* | Within site | 0.382 |
| *Enkianthus campanulatus–Pinus parviflora* | Within site | 0.559 |
| *Enkianthus campanulatus–Rhododendron spp.* | Within site | 0.065 |
| *Enkianthus campanulatus–Vaccinium smallii* | Within site | 0.559 |
| *Eubotryoides grayana–Pinus parviflora* | Within site | 0.021 |
| *Eubotryoides grayana–Rhododendron spp.* | Within site | 0.559 |
| *Eubotryoides grayana–Vaccinium smallii* | Within site | 0.021 |
| *Pinus parviflora–Rhododendron spp.* | Within site | 0.012 |
| *Pinus parviflora–Vaccinium smallii* | Within site | 0.915 |
| *Rhododendron spp.–Vaccinium smallii* | Within site | 0.012 |

(B)

| Host comparison | Permutation restriction | FDR |
|---|---|---|
| *Betula ermanii–Enkianthus campanulatus* | Within site | 0.904 |
| *Betula ermanii–Eubotryoides grayana* | Within site | 0.156 |
| *Betula ermanii–Pinus parviflora* | Within site | 0.904 |
| *Betula ermanii–Rhododendron spp.* | Within site | 0.044 |

| **Host comparison** | **Permutation restriction** | **FDR** |
|---|---|---|
| *Betula ermanii–Vaccinium smallii* | Within site | 0.904 |
| *Enkianthus campanulatus–Eubotryoides grayana* | Within site | 0.038 |
| *Enkianthus campanulatus–Pinus parviflora* | Within site | 0.746 |
| *Enkianthus campanulatus–Rhododendron spp.* | Within site | 0.002 |
| *Enkianthus campanulatus–Vaccinium smallii* | Within site | 0.970 |
| *Eubotryoides grayana–Pinus parviflora* | Within site | 0.146 |
| *Eubotryoides grayana–Rhododendron spp.* | Within site | 0.904 |
| *Eubotryoides grayana–Vaccinium smallii* | Within site | 0.044 |
| *Pinus parviflora–Rhododendron spp.* | Within site | 0.077 |
| *Pinus parviflora–Vaccinium smallii* | Within site | 0.718 |
| *Rhododendron spp.–Vaccinium smallii* | Within site | 0.025 |

**Table S8. Results of PERMANOVA testing for differences in root- and soil-associated community composition across habitats and sample types at representative sampling points.** PERMANOVA was performed separately for (A) prokaryotic and (B) fungal communities based on Sørensen dissimilarity. Habitat differences were tested separately within the root subset and soil samples, with permutations restricted within study sites. Differences between root and soil samples were tested separately within the solfatara field and forest edge, with permutations restricted within sampling points. The table reports degrees of freedom (*df*), sums of squares (SS), proportion of variance explained ($R^2$), *F* statistics (*F*), and *P* values adjusted for multiple comparisons using the Benjamini–Hochberg false discovery rate (FDR) method. $R^2$ represents the proportion of variation explained by each tested factor.

(A)

| Test | Permutation restriction | Factor | *df* | SS | $R^2$ | *F* | FDR |
|---|---|---|---|---|---|---|---|
| Habitat within root subset | Within site | Habitat | 1 | 0.836 | 0.081 | 3.358 | < 0.001 |
| | | Residual | 38 | 9.461 | 0.919 | | |
| | | Total | 39 | 10.297 | 1.000 | | |
| Habitat within soil | | Habitat | 1 | 0.408 | 0.163 | 3.499 | < 0.001 |
| | | Residual | 18 | 2.098 | 0.837 | | |
| | | Total | 19 | 2.506 | 1.000 | | |
| Sample type within solfatara field | Within sampling point | Sample type | 1 | 2.057 | 0.279 | 10.826 | < 0.001 |
| | | Residual | 28 | 5.321 | 0.721 | | |
| | | Total | 29 | 7.379 | 1.000 | | |
| Sample type within forest edge | | Sample type | 1 | 0.741 | 0.106 | 3.325 | < 0.001 |
| | | Residual | 28 | 6.237 | 0.894 | | |
| | | Total | 29 | 6.978 | 1.000 | | |

(B)

| Test | Permutation restriction | Factor | *df* | SS | $R^2$ | *F* | FDR |
|---|---|---|---|---|---|---|---|
| Habitat within root subset | Within site | Habitat | 1 | 2.058 | 0.127 | 5.684 | < 0.001 |
| | | Residual | 39 | 14.121 | 0.873 | | |
| | | Total | 40 | 16.179 | 1.000 | | |
| Habitat within soil | | Habitat | 1 | 1.649 | 0.291 | 7.385 | < 0.001 |

| **Test** | **Permutation restriction** | **Factor** | ***df*** | **SS** | $R^2$ | ***F*** | **FDR** |
|---|---|---|---|---|---|---|---|
| | | Residual | 18 | 4.019 | 0.709 | | |
| | | Total | 19 | 5.669 | 1.000 | | |
| Sample type within solfatara field | Within sampling point | Sample type | 1 | 2.042 | 0.188 | 6.496 | < 0.001 |
| | | Residual | 28 | 8.801 | 0.812 | | |
| | | Total | 29 | 10.843 | 1.000 | | |
| Sample type within forest edge | | Sample type | 1 | 1.280 | 0.121 | 3.975 | < 0.001 |
| | | Residual | 29 | 9.340 | 0.879 | | |
| | | Total | 30 | 10.620 | 1.000 | | |

**Table S9. Variation partitioning of prokaryotic and fungal community composition between habitat and sample type.**

Variation partitioning was performed based on Sørensen dissimilarity for (A) prokaryotic and (B) fungal communities to quantify the fractions of variation attributable to habitat and sample type (root or soil). Fractions are expressed as adjusted $R^2$ values, accounting for model complexity. "Habitat + Sample type" represents the variation explained by the full model containing both variables. "Habitat (unique)" and "Sample type (unique)" represent the fractions uniquely attributable to each variable after accounting for the other, whereas "Shared (Habitat ∩ Sample type)" represents their joint fraction. "Habitat (total)" and "Sample type (total)" represent the variation explained by each variable without accounting for the other. Negative shared fractions can arise from the adjustment of $R^2$ and should be interpreted as indicating essentially no shared explained variation rather than as a negative amount of variance.

(A)

| **Partition** | **Adjusted $R^2$** | ***df*** |
|---|---|---|
| Habitat (total) | 0.038 | 1 |
| Sample type (total) | 0.142 | 1 |
| Habitat + Sample type | 0.183 | 2 |
| Habitat (unique) | 0.041 | 1 |
| Sample type (unique) | 0.145 | 1 |
| Shared (Habitat ∩ Sample type) | -0.003 | 0 |
| Residual | 0.817 | |

(B)

| **Partition** | **Adjusted $R^2$** | ***df*** |
|---|---|---|
| Habitat (total) | 0.098 | 1 |
| Sample type (total) | 0.082 | 1 |
| Habitat + Sample type | 0.183 | 2 |
| Habitat (unique) | 0.101 | 1 |
| Sample type (unique) | 0.085 | 1 |
| Shared (Habitat ∩ Sample type) | -0.003 | 0 |
| Residual | 0.817 | |

**Table S10. Results of PERMDISP testing for differences in within-group dispersion of root- and soil-associated community composition across habitats and sample types.**

PERMDISP was performed separately for (A) prokaryotic and (B) fungal communities based on Sørensen dissimilarity. Habitat differences were tested separately within the root subset and soil samples, with permutations restricted within study sites. Differences between root and soil samples were tested separately within the solfatara field and forest edge, with permutations restricted within sampling points. Significant results indicate heterogeneity in multivariate dispersion, that is, differences in within-group variation in community composition. The table reports degrees of freedom (*df*), sums of squares (SS), mean squares (MS), *F* statistics (*F*), and *P* values adjusted for multiple comparisons using the Benjamini–Hochberg false discovery rate (FDR) method.

(A)

| **Test** | **Permutation restriction** | **Factor** | ***df*** | **SS** | **MS** | ***F*** | **FDR** |
|---|---|---|---|---|---|---|---|
| Habitat within root subset | Within site | Habitat | 1 | 0.007 | 0.007 | 0.482 | 0.668 |
| | | Residual | 38 | 0.580 | 0.015 | | |
| Habitat within soil | | Habitat | 1 | 0.001 | 0.001 | 0.187 | 0.668 |
| | | Residual | 18 | 0.067 | 0.004 | | |
| Sample type within solfatara field | Within sampling point | Sample type | 1 | 0.133 | 0.133 | 35.488 | < 0.001 |
| | | Residual | 28 | 0.105 | 0.004 | | |
| Sample type within forest edge | | Sample type | 1 | 0.163 | 0.163 | 8.394 | < 0.001 |
| | | Residual | 28 | 0.544 | 0.019 | | |

(B)

| **Test** | **Permutation restriction** | **Factor** | ***df*** | **SS** | **MS** | ***F*** | **FDR** |
|---|---|---|---|---|---|---|---|
| Habitat within root subset | Within site | Habitat | 1 | 0.001 | 0.001 | 0.067 | 0.760 |
| | | Residual | 39 | 0.494 | 0.013 | | |
| Habitat within soil | | Habitat | 1 | 0.009 | 0.009 | 0.876 | 0.735 |
| | | Residual | 18 | 0.185 | 0.010 | | |
| Sample type within solfatara field | Within sampling point | Sample type | 1 | 0.160 | 0.160 | 14.725 | < 0.001 |
| | | Residual | 28 | 0.305 | 0.011 | | |
| Sample type within forest edge | | Sample type | 1 | 0.072 | 0.072 | 5.696 | < 0.001 |
| | | Residual | 29 | 0.368 | 0.013 | | |
